# Lipid-directed covalent fluorescent labeling of plasma membranes for long-term imaging, barcoding and manipulation of cells

**DOI:** 10.1101/2024.07.28.605476

**Authors:** Nathan Aknine, Remi Pelletier, Andrey S. Klymchenko

## Abstract

Fluorescent probes for cell plasma membrane (PM) are generally based on amphiphilic anchors that incorporate non-covalently into biomembranes. Therefore, they are not compatible with fixation and permeabilization, presence of serum, or cell co-culture because of their exchange with the medium. Here, we report a concept of lipid-directed covalent labeling of PM, which exploits transient binding to lipid membrane surface generating high local dye concentration, thus favoring covalent ligation to random proximal membrane proteins. This concept yielded a class of fluorescent probes for PM (MemGraft), where a cyanine dye (Cy3 and Cy5) bears at its two ends low-affinity membrane anchor and reactive group: an activated ester or a maleimide. We found that MemGraft probes with these reactive groups provide efficient PM labelling, in contrast to a series of control compounds, including commercial Cy3-based labels of amino and thiol groups, revealing the crucial role of the membrane anchor combined with high reactivity of activated ester and a maleimide groups. In contrast to conventional PM probes, based on non-covalent interactions, MemGraft labelling approach is compatible with cell fixation, permeabilization, trypsinization and presence of serum. The latter allows long-term cell tracking and video imaging of cell PM dynamics without signs of phototoxicity. The covalent strategy also enables staining and long-term tracking of co-cultured cells labelled in different colors without probes exchange. Moreover, combination of different ratios of MemGraft-Cy3 and MemGraft-Cy5 probes enabled long-term cell barcoding in at least 5 color codes, important for tracking and visualizing multiple cells populations. Ultimately, we found that MemGraft strategy enables efficient biotinylation of cell surface, opening the path to cell surface engineering and cell manipulation.

## Introduction

Cell plasma membrane (PM) is an indispensable frontier at the cell surface. It is not a simple barrier between the cell interior and the environment, but a highly organized two-dimensional structure with specially organized lipids and proteins, which plays essential roles in the signaling, transport of nutrients, cell-cell recognition, motility, etc. ^1-3^ Therefore, the development of fluorescent tools to image and probe cell plasma membranes is currently a subject of intensive research.^4-7^ So far, a large variety of fluorescent membrane probes have been developed. Some operate like basic membrane markers, which include for instance cyanine derivatives, like DiO, DiI, PKH, and more advanced analogs, such as MemBright.^8,9^ Other probes are designed for sensing local membrane properties, including local polarity, ^10-12^ viscosity, ^13,14^ and tension^15-17^ that help to understand lipid organization, or detect local pH, ^18^ ions, ^19^ specific lipids^20,21^ or membrane proteins. ^22,23^ However, most of those examples are based on lipophilic or amphiphilic lipid anchors or protein ligands, which interact non-covalently with cell plasma membranes. The non-covalent labeling of PM implies that the probes could exchange with the environment, in particular membranes of other cells or serum proteins.

Moreover, after fixation and especially permeabilization, used in a variety of staining protocols especially immunofluorescence,^24,25^ these probes are generally washed away from the cell surface, because membrane integrity is compromised and most of the lipids and lipid probes are removed by the detergent. These issues make those probes also incompatible with expansion microscopy,^26-28^ tissue clearing ^29-31^ and correlative light and electron microscopy (CLEM).^32,33^ Moreover, fast lateral diffusion of PM probes makes them poorly compatible with super-resolution imaging, in particular single-molecule localization microscopy.^34-38^ A promising solution is the covalent grafting of fluorescent dyes to proteins, which are compatible with both fixation and permeabilization protocols. The efficient labelling approaches include the use of potent tags, such as SnapTag,^39,40^ HaloTag,^41,42^ ClipTag, ^43^, etc, but they require genetic modification of the cell. Direct labeling of native proteins on the cell surface could be done for instance by activated esters of N-hydroxysuccinimide (NHS). However, this approach is not so convenient and efficient, because this labeling reaction requires high concentrations of the labeling agent (∼5 μM).^44^ Therefore, we hypothesized that the labeling could be accelerated by designing a reactive probe capable of first binding to the cell surface non-covalently. Then, the high local concentration of the probe would favor the fast labeling of neighboring proteins. In this respect, one should mention affinity-based protein labelling,^45,46^ where an affinity ligand that binds a specific protein site favors covalent grafting to a proximal reactive group of the protein. It was successfully applied as covalent inhibitors in drug discovery,^47,48^ as well as for site specific labelling of antibodies^45,49,50^ and specific labelling of membrane receptors.^51-54^ However, these are approaches for selective protein labelling, which would target only a limited fraction of specific proteins, inefficient for bright staining of PM. Therefore, we considered a different strategy where the designed probe would label non-selectively a large variety of membrane proteins. For this purpose, we considered exploiting non-covalent interaction with lipids. The abundance of lipids on the PM would ensure that membrane bound probes would label randomly any available membrane protein, which could provide efficient high density covalent plasma membrane labeling. Although the targeting of PM is usually done with high-affinity membrane probes,^4-7^ this is not useful in this case because it would be difficult to remove non-reacted probes after the labeling. In this respect, it would be interesting to explore low-affinity membrane targeting groups, which could reversibly bind to cell plasma membranes. Recently, we introduced low-affinity amphiphilic anchors for plasma membrane targeting composed of butyl chain and charged sulfonate head group. ^11,38^ Reversible low-affinity binding of probes based on solvatochromic Nile Red^11^ and fluorogenic dimer^38^ enabled PAINT-type super-resolution imaging of plasma membranes. However, it remains unexplored whether this reversible binding could favor a reaction between activated ester or maleimide group and membrane proteins, which would enable efficient covalent grafting of fluorescent probes to plasma membranes.

In the present work, we designed a series of cyanine derivatives bearing a low-affinity membrane targeting anchor group in combination with a reactive group for conjugation. We found that the combination of low-affinity anchor with the activated ester or maleimide group provides strong labeling of the cell surface for both Cy3 and Cy5 analogs, which was not the case for non-activated carboxylate control analogs. Moreover, the control compound with a truncated anchor (without butyl group) does not show plasma membrane labeling, which highlights that the membrane protein labeling is catalyzed by probe-lipid interactions. Importantly, obtained plasma membrane labeling showed resistance to the presence of serum as well as to cell fixation and permeabilization in contrast to a reference plasma membrane probe. The MemGraft strategy also enable long-term cell tracking and multi-color barcoding of multiple co-cultured cell populations. The covalent strategy also allowed design MemGraft probe for cell biotinylation and further cell manipulation with magnetic beads. The developed probe will find multiple applications in cellular imaging, cell surface engineering and cell manipulation, where robust grafting to the cell surface is needed.

## Results and discussion

### Design and synthesis

The design of the probes is based on a cyanine fluorophore, bearing on one end low-affinity membrane anchor composed of a butyl chain and sulfonate group, while the other end of cyanine bears an activated ester (Figure 1A, B). In our hypothesis, the low-affinity membrane anchor would provide transient binding to a lipid membrane, as we showed for Nile Red^11^ and cyanine dimers, ^38^ which is expected to ensure a high local concentration of the probe at the membrane level (vs buffer) in close proximity to membrane proteins (Figure 1). The latter should favor covalent grafting of the active ester with amino groups of membrane proteins, needed for covalent labelling of plasma membranes (Figure 1). The synthesis was based on our previous report,^38^ where corresponding cyanine (Cy3 or Cy5) dicarboxylate, was first monofunctionalized with the anchor (3-(butylammonio)propane-1-sulfonate) and then, after purification, the second carboxylate was converted into activated NHS ester using TSTU (Figure S1). These compounds will be called here MemGraft-Cy3 and MemGraft-Cy5 probes, respectively (Figure 1). To understand the role of the anchor group, we synthesized Cy3 analogs without butyl chain (dye **1**) and with octyl chain (dye 2). To this end, the Cy3 dicarboxylate was first monofunctionalized by 3-aminopropane-1-sulfonate or 3-(octylammonio)propane-1-sulfonate and the second carboxylate was converted into NHS ester using TSTU to yield dyes **1** and **2**, respectively. Then, to explore the importance of the activated ester, we also synthesized MemGraft-Cy3 analogs, bearing fluorophenyl activated esters of increasing reactivity (Figure 1): 2-fluorophenyl (dye **3**) and 2,6-diflurophenyl (dye **4**), 2,3,5,6-tetraflurophenyl (dye **5**) and 2,3,4,5,6-pentaflurophenyl (dye **6**), which are also expected to react with amino groups of proteins.^55^ The non-activated control compounds for MemGraft-Cy3 and MemGraft-Cy5 probes were intermediate carboxylic acid analogues: dyes **7** and **8**, respectively. Moreover, to expand the approach to other type of bioconjugation, we replaced an activated ester with a maleimide group (MemGraft-Cy3M), which is expected to react with thiol groups of membrane proteins.

**Figure 1.**
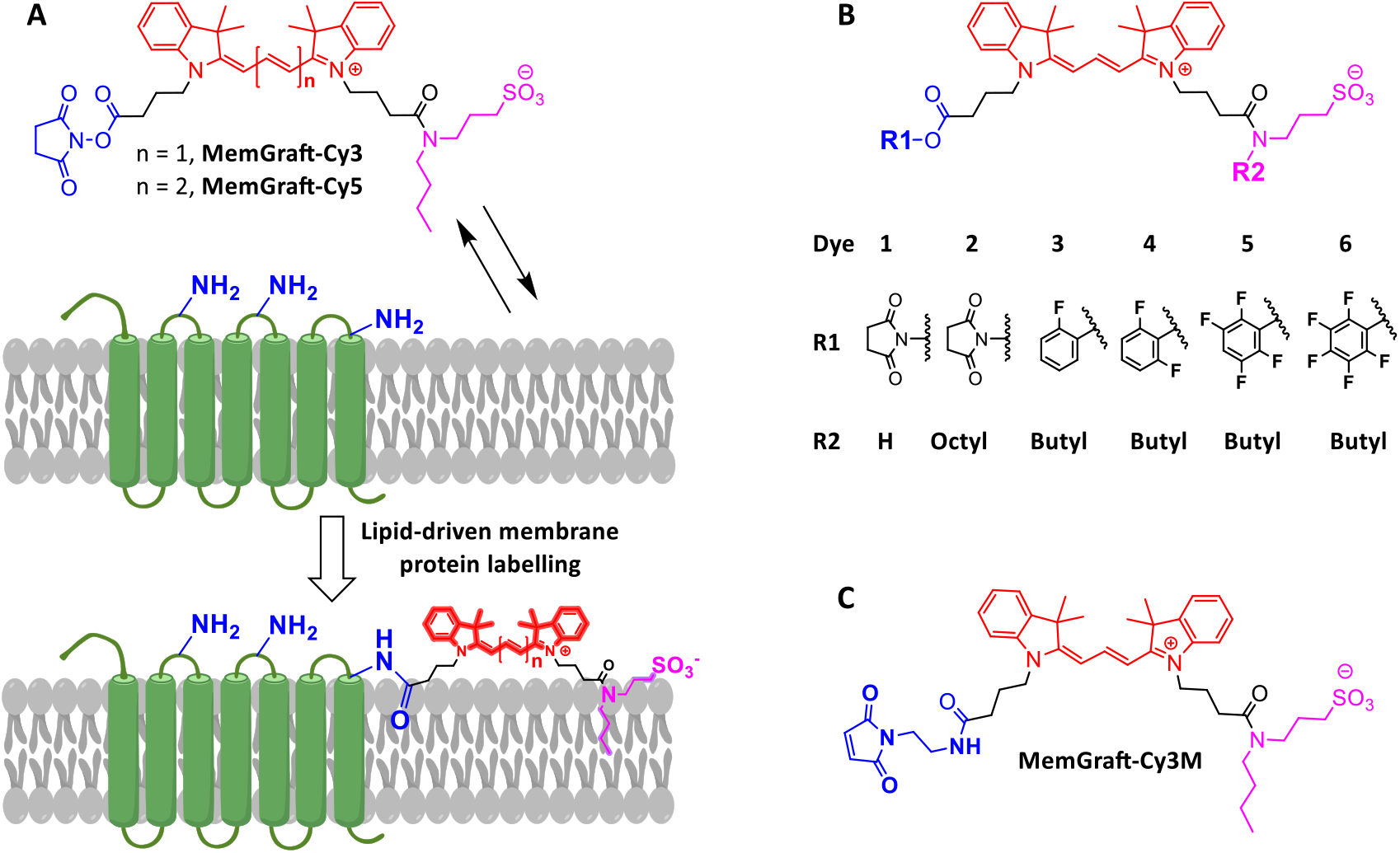
MemGraft probes and the concept of lipid driven labelling of membrane proteins. (A) Chemical structures of MemGraft (A) and control (B) probes.

### Labelling of cell plasma membranes

First, we characterized the most reactive activated ester (NHS) derivatives. The Cy3 and Cy5 probes (MemGraft-Cy3, and MemGraft-Cy5, respectively) were incubated with live cells for 5 min and then imaged after washing. A clear cell surface fluorescence was observed for both Cy3- and Cy5-based probes in U87 cells (Figure 2). Imaging experiments on another cell line (HeLa) with MemGraft-Cy3 showed similar cell surface labelling (Figure S2). The staining profiles of MemGraft-Cy3 and MemGraft-Cy5 were nearly identical to that of a reference probe MemBright-488 (Figure S3), suggesting efficient PM targeting. At the same labelling conditions, control analogs without activated carboxylate (dyes **7** and **8**) showed practically no membrane staining (Figure 2). The latter suggests that, as designed, these control probes exhibit low affinity to plasma membranes and therefore can be completely removed by washing. Therefore, the signal from the activated probes at the plasma membranes corresponds exclusively to the covalently grafted probes. To understand the role of hydrophobic interactions of the probes in the membrane protein labeling, we varied the alkyl chain of the anchor of MemGraft-Cy3. Thus, we designed an analogue without any alkyl group (dye **1**) and with an octyl group (dye **2**), which are expected to strongly alder their affinity to lipid membranes compared to the patent MemGraft-Cy3. Remarkably, the control probe **1** showed mainly intracellular fluorescence without any significant plasma membrane labelling, in contrast to MemGraft-Cy3 (Figure 3). This dramatic change in the labelling profile caused by the removal of the butyl group indicates that the latter plays a crucial role in PM labeling. On the other hand, dye **2** with octyl chain showed good PM labelling, in contrast to its non-activated acid analogue **10** (Figure 3). Indeed, the butyl or octyl groups should favor transient binding and certain accumulation of the probe on the lipid membrane surface.^11^ This is expected to ensure a high local concentration of the MemGraft probes that would favor the covalent grafting to membrane proteins (Figure 1). On the other hand, the analogue without the butyl group cannot stay sufficiently long at the cell surface and show some capacity to penetrate PM, which can explain the lack of PM labeling and the significant intracellular signal. A similar difference between MemGraft-Cy3 and the control dye **1** without butyl group was observed at a lower concentration (0.1 μM, Figure S4), which additionally shows that the MemGraft probe can efficiently label PM despite this low dye concentration.

**Figure 2.**
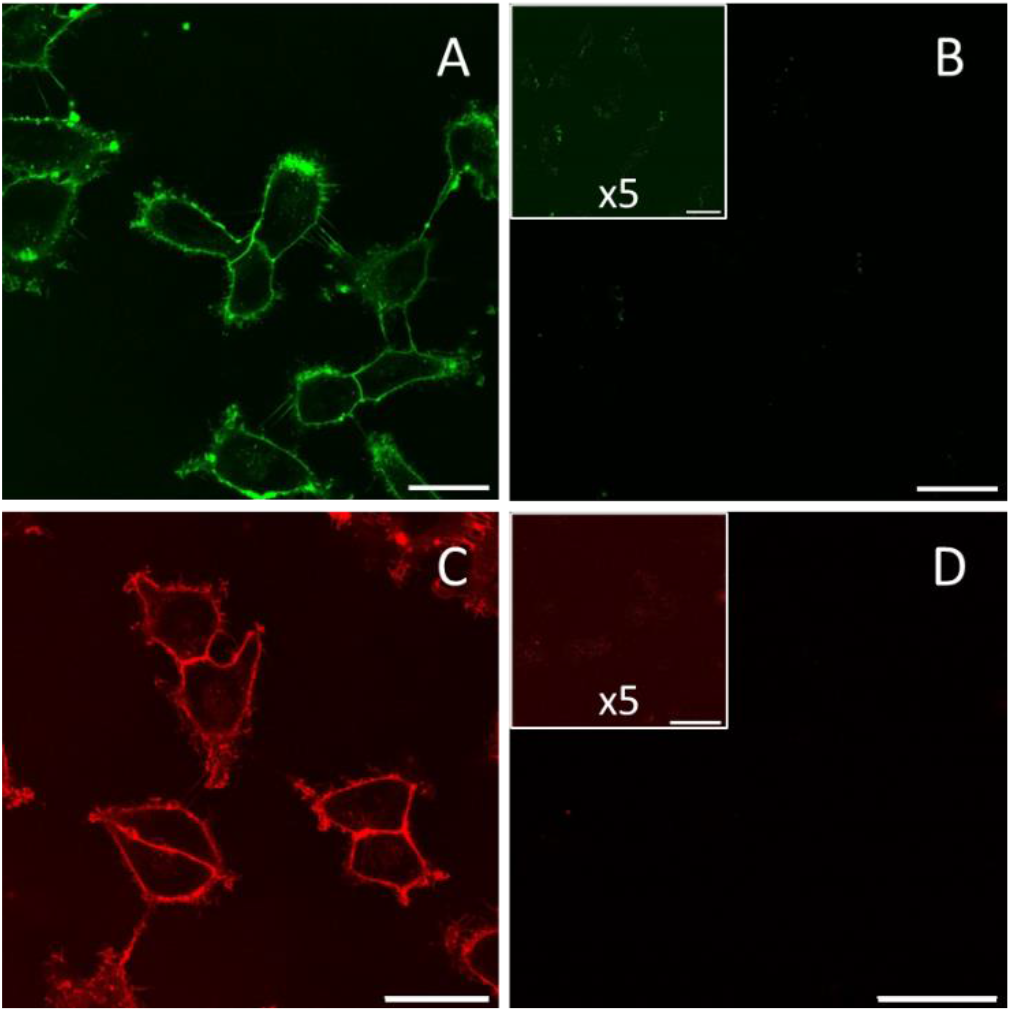
Fluorescence labeling of plasma membrane using MemGraft-Cy3 (A) and MemGraft-Cy5 (C) dyes and their non-activated analogues **7** (B) and **8** (D). Confocal fluorescence microscopy of U87 cells incubated for 5 min with the dyes at 1 μM concentration. Scale bar: 30μm. The insets of B and D are the same images with 5-fold multiplied intensity to show the lack of labeling.

**Figure 3.**
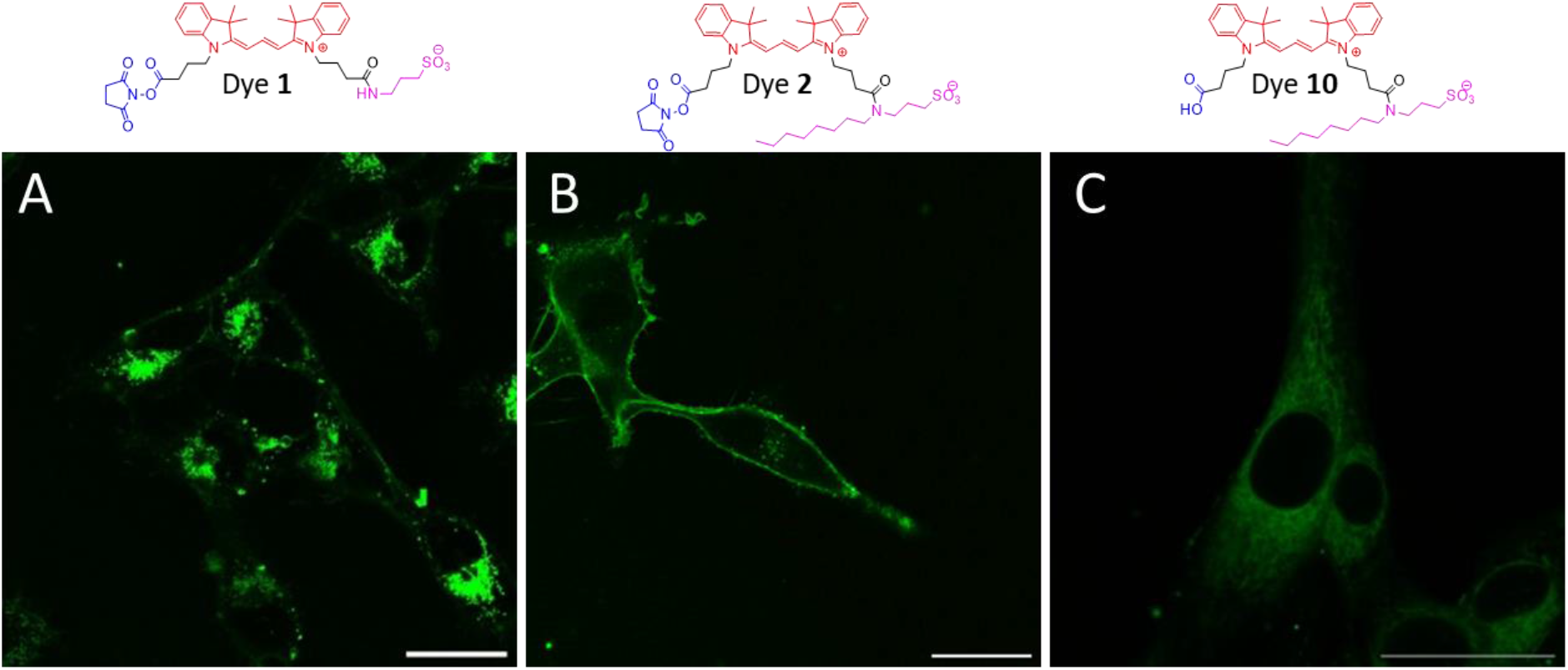
The role of membrane anchor in analogues of MemGraft-Cy3. Confocal fluorescence microscopy of U87 cells incubated with dyes **1** (A), **2** (B) and **10** (C). Dye concentration was 1 μM. Scale bar: 30 μm.

Then, we tested control dyes **3**-**6**, which bear activated esters for amino group labelling of varied reactivity. However, they did not provide efficient plasma membrane labelling, but rather significant intracellular staining (Figure S5). Nevertheless, we observed an interesting tendency. While compounds **3** and **4** with lower reactivity towards amino groups showed exclusively intracellular staining, more reactive dyes **5** and **6** exhibited a clearly seen cell contour (Figure S5). The membrane staining was the best seen for the most reactive dye **6** in this series, where membrane extensions could be well identified, although the intracellular fluorescence remained strong (Figure S5). These results indicate that reactivity of the active ester is crucial for an efficient plasma membrane labelling. We can conclude that, in contrast to NHS esters, none of the fluorophenyl esters is reactive enough, as the internalization goes faster than the protein labelling. Moreover, fluorophenyl esters are much more hydrophobic than the NHS one, which could additionally favor internalization by crossing the plasma membrane.

We also compared our MemGraft probes with commercial reference dye disulfo-Cy3-NHS, which is routinely used for amino-group labelling of proteins. Fluorescence microscopy of cells labelled with disulfo-Cy3-NHS at the same experimental conditions showed a significantly smaller signal at the cell surface compared to MemGraft-Cy3 (Figure 4A,B). This result confirms that the presence of membrane anchor significantly improves membrane labelling, because it favors higher membrane concentrations of the dye that can catalyze the reaction with the amino groups of membrane proteins. To further shed light on the nature of the labelling, the amino residues on the cell surface were blocked by treatment with Sulfo-NHS-acetate on cells.^56^ This is a highly polar agent, which is expected to acetylate only water-exposed amino-group residues at the cell surface. After the treatment, the cells were stained with MemGraft-Cy3 or disulfo-Cy3-NHS and the fluorescence intensity of the cells was compared to those without the blocking agent (Figure S6). Remarkably, the blocking by Sulfo-NHS-acetate nearly totally inhibited cell membrane labelling disulfo-Cy3-NHS with only 8% residual fluorescence signal, while the staining by MemGraft-Cy3 was decreased to a small extent (60% of the residual fluorescence signal). This proves that the nature of the MemGraft labelling is different from classical polar NHS-ester reagents, like disulfo-Cy3-NHS. Due to the presence of the low-affinity membrane anchor, MemGraft-Cy3 probably labels amino groups of proteins in close contact with the lipid bilayer that cannot be accessed by Sulfo-NHS-acetate (Figure S6), while commercial disulfo-Cy3-NHS labels exposed amino groups assessable for Sulfo-NHS-acetate. Then, we also tested maleimide analogue MemGraft-Cy3M and compared it to a commercial thiol label based on Cy3, disulfo-Cy3-MAL. Remarkably, MemGraft-Cy3M showed clear labelling of PM, while disulfo-Cy3-MAL failed to label PM (Figure 4C,D). This result confirms the crucial role of low-affinity membrane anchor for covalent PM labeling. Disulfo-Cy3-MAL did not label PM at all probably because thiol groups of membrane proteins are not accessible for this highly polar label, while MemGraft-Cy3M, owing to its membrane binding, can reach and react with membrane-shielded thiol groups.

**Figure 4.**
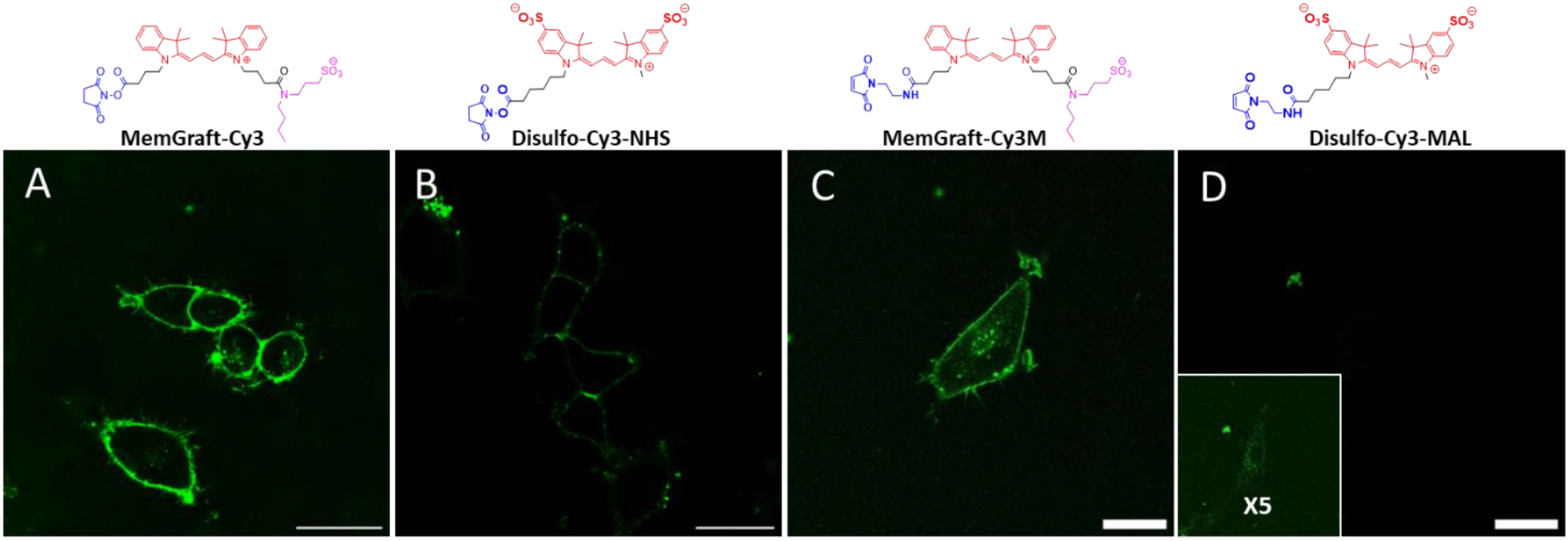
Comparison of MemGraft-Cy3 and MemGraft-Cy3M with commercial labels of close structure. (A, B) Confocal images of U87 cells incubated with MemGraft-Cy3 (C) and disulfo-Cy3-NHS (D) at 100 nM concentration. Scale bar: 30 μm. (C, D) Confocal images of U87 cells incubated with MemGraft-Cy3M (C) and disulfo-Cy3-MAL (D) at 500 nM concentration. Scale bar: 30 μm.

The labeling density of MemGraft-Cy3 was evaluated by flow cytometry in comparison with the commercial label disulfo-Cy3-NHS. A population of 25000-60000 single cells was first identified by SSC and FSC analysis and the fluorescence signals from single cells were then evaluated for both covalent probes at the same concentrations (0.1 or 1 μM). For both studied concentrations, the fluorescence mean intensity of MemGraft-Cy3 probe was 3 times higher than that of disulfo-Cy3-NHS (Figure 5). This result suggests a higher density of labeling for MemGraft-Cy3 compared to its analog without membrane anchor, in line with microscopy data (Figure 4A,B). Thus, the cytometry data confirmed the superiority of MemGraft approach compared to direct labelling of amino groups by a commercial dye NHS-ester. It highlights the importance of low-affinity membrane anchor in MemGraft that allows the pre-organization of the reactive probe close to amino groups of the membrane proteins, required for efficient membrane labelling. It enables relatively high labelling density for rather low probe concentrations from 0.1 to 1 μM.

**Figure 5.**
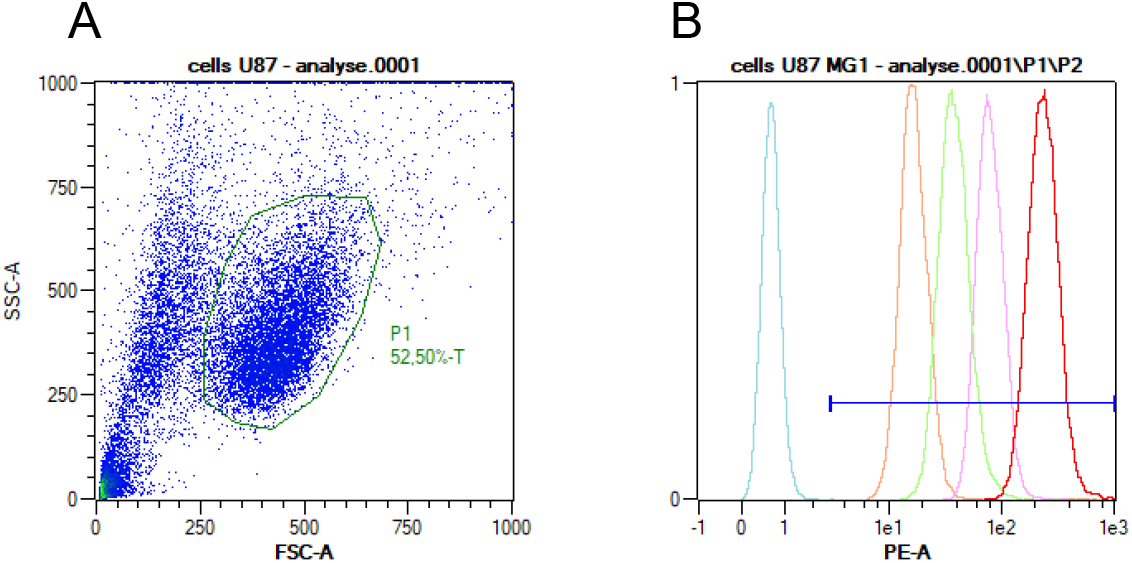
(a) Representative flow cytometry density plot of U87 cells showing side scattered light (granularity) vs. forward scattered light (size of cells). The region P1 is established from non-treated cells. Cells doublets are abstracted by filtrating counts out of linearity in FSC-A vs FSC-H plot. (b) Normalized flow cytometry histograms showing cell counts vs. fluorescence intensity of not stained cells (blue), cells stained with disulfo-Cy3-NHS 0.1 μM (orange) and 1 μM (pink) or MemGraft-Cy3 at 0.1 μM (green) and 1 μM (red).

The conjugation between MemGraft probe and a model membrane containing surface amino groups was realized to confirm the chemistry of the labeling. Large Unilamellar vesicles (LUVs) were prepared with a lipid composition of DOPC/DOPE (1:1). The formed liposomes present exofacial reactive amino groups on their outer surface, which are expected to react with MemGraft. The liposomes were incubated with MemGraft-Cy3 probe, followed by purification by size-exclusion column chromatography and further mass spectrometry. We found a mass-to-charge ratio at 1403.58, corresponding to the conjugate of MemGraft-Cy3 with DOPE (Figure S7). This result confirms the capacity of MemGraft-Cy3 to form covalent bonds with amino groups at the membrane surface, which confirms the covalent nature of the labelling. However, on the cell surface, it is most likely the probe reacts with the amino-groups of membrane proteins, as PE is lacking at the outer leaflet of healthy cell plasma membrane. To verify this point, we performed SDS-page on cytosolic and membraneprotein extracts. The obtained results suggest that a large range of proteins was labelled by MemGraft-Cy3, as it could be seen by a contentious signal over multiple molecular masses from 20 to 250 kDa (Figure S8). In case of MemGraft-Cy3M, the signal was much weaker, but still visible for proteins of different molecular weight, with predominance of one fraction around 50 kDa. In contrast, cytosolic proteins did not show any signal for both MemGraft probes, confirming that the labelling is specific to membrane proteins only. Thus, as expected, chemically reactive MemGraft probes label non-selectively membrane proteins, which is essential for the efficient labelling of cell plasma membranes.

### Resistance to serum

Next, we evaluated the capacity of the new probes to remain on the membrane surface despite the presence of full growth medium containing serum, which is essential for cell growth. The MemGraft-Cy3 probe was compared to a commercial reference probe (MemBright-488) that binds non-covalently to the plasma membrane.^9^ After 5 min incubation in 10% FBS, the membrane staining by MemGraft-Cy3 probe remains similar to that of reference probe MemBright-488 (Figure 6). However, after longer incubation in the presence of serum, the PM signal of MemBright-488 progressively declined and became poorly detectable after 45 min. The remaining signal appeared mainly as perinuclear dotted fluorescence, corresponding probably to endosomes. In contrast, in the presence of 10% FBS, the membrane staining by MemGraft-Cy3 remained similar to that without FBS for the whole observation period, even though a small fraction of intracellular dots could be also observed after 45 min, clearly due to the endocytosis (Figure 6). Thus, we can conclude that FBS can detach Membright-488 probe from the cell surface, whereas the remaining fraction of the probe is internalized by endocytosis. On the other hand, MemGraft-Cy3 probe resisted well to FBS, so only endocytosis process affected to some extent the imaging contrast over time. Thus, the MemGraft probe enables imaging of PM in live cells in the presence of serum, which was not possible before with conventional membrane markers. This is important for longitudinal studies of cell behavior without cell starvation.

**Figure 6.**
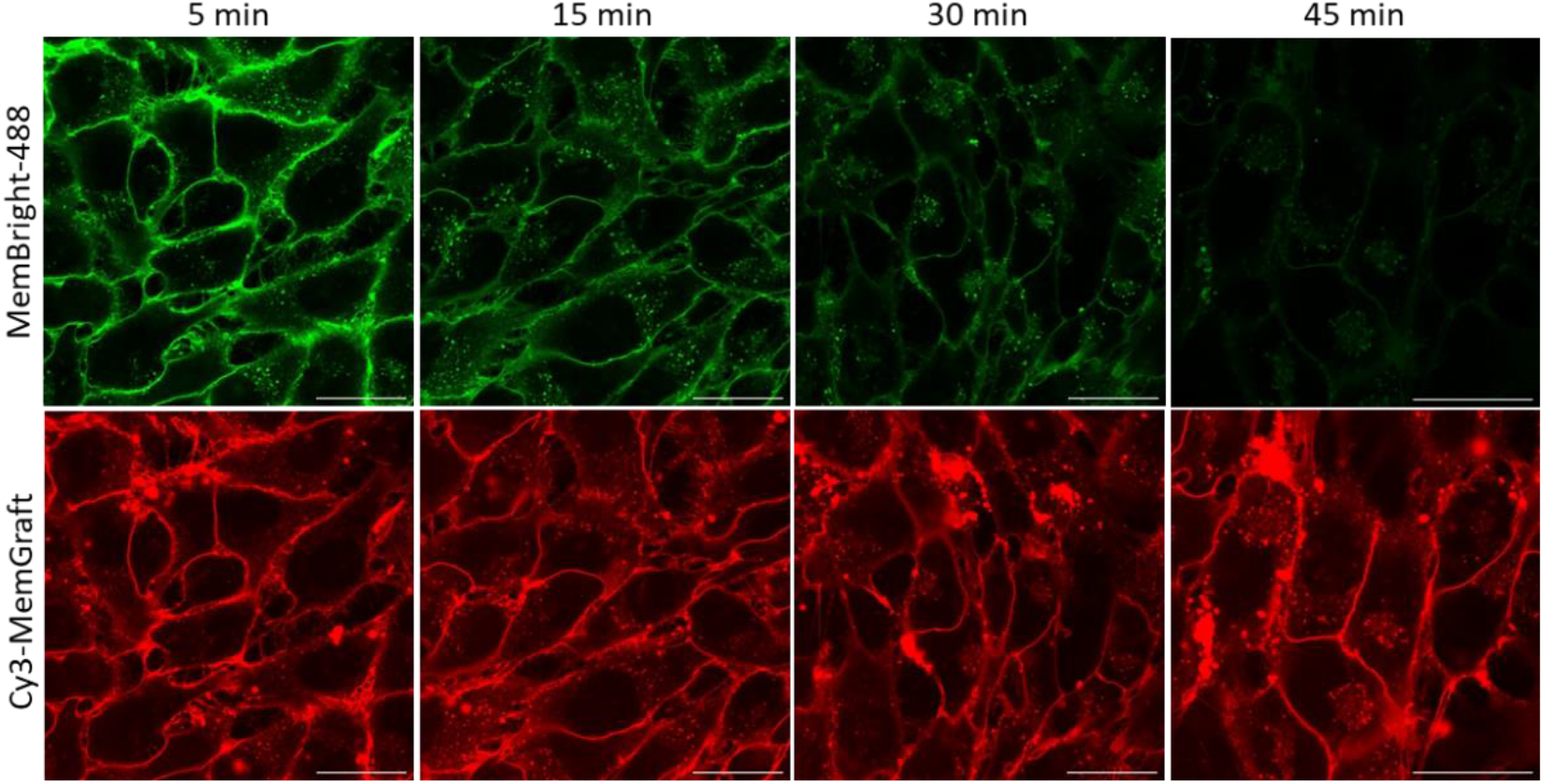
Resistance of new probes to the presence of fetal bovine serum (FBS) in the growth medium. Confocal fluorescence microscopy of U87 cells incubated with MemGraft-Cy3 (red) and reference MemBright-488 probe (green) with 10% FBS. Dye concentration was 1 μM. Scale bar: 30 μm.

Therefore, the U87 cells were stained with MemGraft-Cy3 and further studied using the long-term imaging in serum-containing medium at 37 °C and 5% CO_2_. Remarkably, we were able to perform continuous video imaging of cells for at least 3h without significant loss of intensity or cell death (Video S1). These results confirmed high photostability and low phototoxicity of MemGraft-Cy3 probe, making it a robust tool for long-term imaging of live cells in native conditions of cell culture medium with serum.

### Resistance to fixation and permeabilization

Even more dramatic effects can be produced on membrane probes during fixation and permeabilization of the cells. Indeed, while lipids and conventional membrane probes cannot be fixed by paraformaldehyde (PFA),^57,58^ they are expected to be washed out during the permeabilization with surfactant.^59^ Therefore, we studied the effect of fixation and permeabilization on the cell staining with MemGraft-Cy3 probe in comparison to the reference MemBright-488. After fixation with PFA, both MemBright and MemGraft probes showed plasma membrane staining with good colocalization (Figure 7). This was expected given that the fixation does not directly disturb lipid bilayers and thus the labeling should be preserved even with non-covalent membrane probes. Second, we performed permeabilization of cells, which is commonly used for immunofluorescence. Under permeabilization conditions using a standard protocol based on Tween-20 surfactant, MemBright-488 probe lost specificity to plasma membranes showing the emission signal all over the cells (Figure 7). In contrast, the labeling profile by MemGraft-Cy3 was affected to a minimal extent, showing clear selective plasma membrane staining (Figure 7). Thus, permeabilization based on the detergent treatment led to the removal of lipid bilayers, which led to the loss of plasma membrane staining by MemBright-488, accompanied by internalization of the probe inside the cells. On the other hand, MemGraft-Cy3 probe was attached covalently to membrane proteins, which were fixed by PFA. Then, the application of detergent affected mainly lipids without a strong effect on the fixed membrane proteins, which explains the resistance of the probe to permeabilization. One should note that the MemGraft probe can also help to verify the integrity of the membrane protein organization after detergent treatment. Indeed, after harsher treatment using Triton X100 at 0.2% concentration for 8 min, the staining profile of the cells by MemGraft-Cy3 was less clear, indicating that this permeabilization protocol can alter protein organization at the cell surface (data not shown).

**Figure 7.**
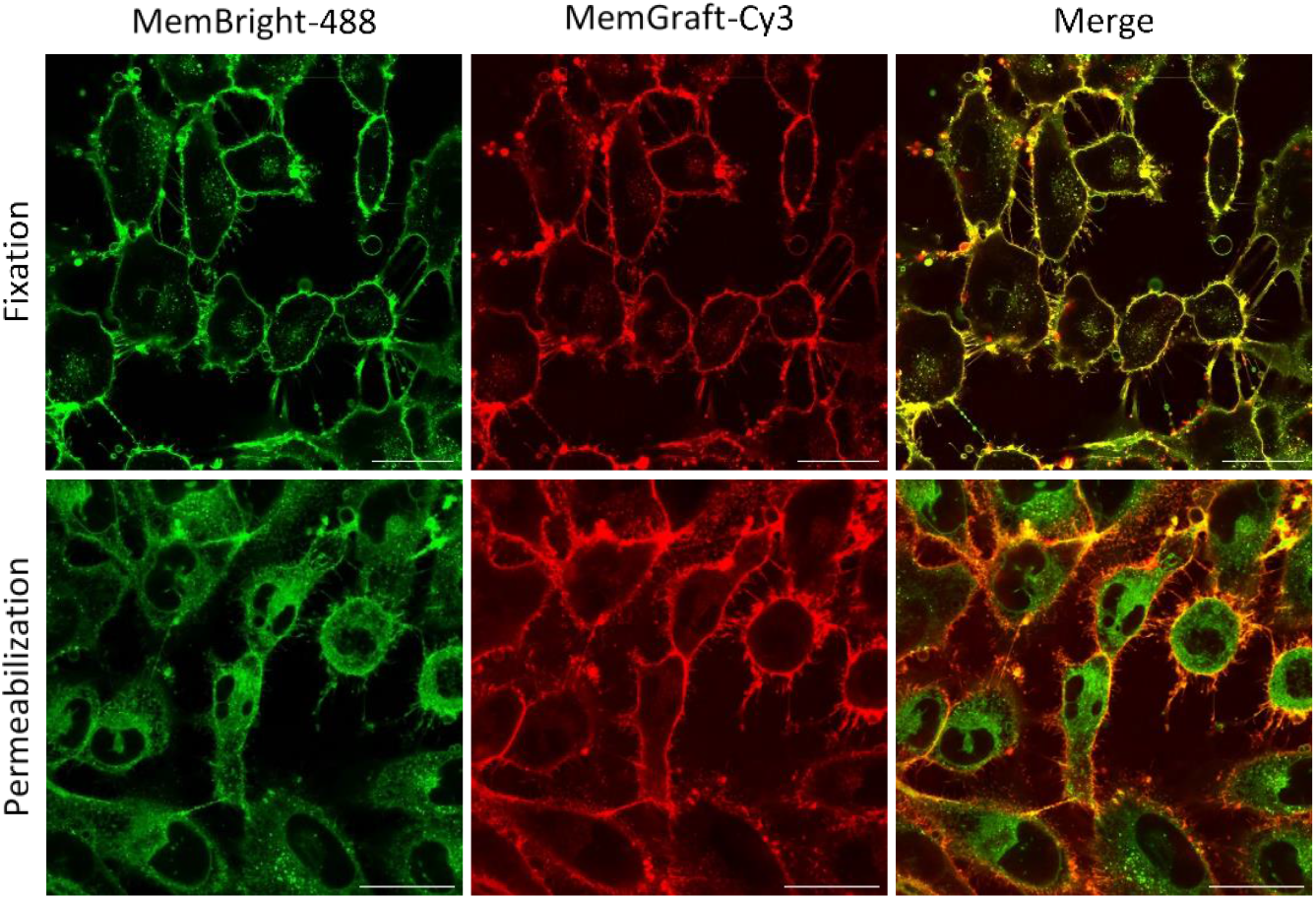
Resistance of MemGraft probe to fixation with permeabilization. Confocal fluorescence microscopy of U87 cells incubated with MemGraft-Cy3 (red) and reference MemBright-488 (green) after fixation with PFA (upper panels) and permeabilization (lower panels) with Tween-20. Dye concentrations: 1 μM for MemGraft-Cy3 and 200 nM for MemBright-488. Scale bar: 30 μm.

### Compatibility with trypsinization and seeding for long-term observation

Trypsinization is a common method to detach the cells from the surface used in cell seeding and other experiments needed in cell transfer from one surface to another with further long-term observation. As MemGraft probes provide covalent labelling to membrane proteins, we studied whether trypsinization could compromise this labelling. To this end, we incubated the cells with the MemGraft-Cy3 and MemBright-488 probes, then after washing detached and seeded them into another microscopy plate. Remarkably, the MemGraft-Cy3 preserved strong signal at the plasma membrane after trypsinization, while MemBright-488 did not (Figure S9). The resistance of MemGraft-Cy3 to trypsinization is surprising because one would expect that cleavage of proteins at the cell surface could lead to the removal of the covalent probe. We could explain this notable resistance to trypsinization by its labelling to amino residues of proteins close to lipid membranes, which are probably not assessable to the enzyme, in line with the lack of accessibility observed for the highly polar acetylation reagent sulfo-NHS-acetate as mentioned above.

### Long-term cell tracking

One of the key requirements for the long-term cell experiments is low cytotoxicity of the label. Therefore, we performed cell viability study for MemBright-Cy5 probe after 24 h incubation (MemBright-Cy3 was not studied to avoid a cross-talk with the MTT assay). The results showed relatively low cytotoxicity of the probe, as >80% of cell viability was observed for concentration range up to 5000 nM (Figure S10). Next, we tested the possibility of using MemGraft for tracking cells and their interactions with other cells in natural growth medium. Classical non-covalent membrane labeling by membrane probes is not suitable for this purpose because serum extracts the probe and the liberated dye could be exchanged between the cells. Two populations of U87 cells were labeled separately by MemGraft-Cy3 and MemGraft-Cy5 and washed. The cells were detached by accutase treatment (milder than trypsinization) and after centrifugation and washing, they were seeded either separately or together in the full-growth medium with FBS. After 5h of the labelling, both probes showed significant labelling of the cell plasma membrane as well as perinuclear dots, corresponding to endosomes (Figure 8, S11). Thus, as expected, some fraction of the probes internalized, while the remaining fraction stayed at the plasma membrane despite the presence of FBS in the medium. Importantly, MemGraft-Cy3 and MemGraft-Cy5 labeled cells incubated separately were observed exclusively in Cy3 and Cy5 channels, respectively, with minimal cross-talk between the channels (Figure 8-A2-B1), which should allow us to distinguish well the signal from the two different probes. The co-incubated cells labeled with two different probes resulted in the observation of two cell populations of different colors (Figure 8-C3-D3). The cells of different colors showed clearly distinctive signals from the individual probes without signs of mixing (Figure 8, S11). It is important to note the individual colors without mixing were observed even for cells in direct contact with each other (Figure 8-C3). The only exception was the direct contact between the two membranes, which appeared in yellow (Figure 8-C3), because the distance between the two plasma membranes was below the resolution of the optical microscopy. This shows that even in the proximity, the probes grafted to the membrane proteins do not exchange with the neighboring cells allowing tracking the interaction of the individual cells. After 29h co-incubation of two differently labeled cell populations, we could still observe very similar staining profiles where plasma membranes labeled with MemGraft probes could be clearly identified. Due to the cell division, the cell density was higher so most of the cells were in direct contact with each other. Nevertheless, the individual cells labeled with two distinct colors of MemGraft-Cy3 and MemGraft-Cy5 probes could still be clearly identified with relatively small color mixing. Some intracellular dots appeared in yellow-orange indicating that cells could internalize membrane proteins of other cells, probably originating from dead cells or exosomes. A better understanding of the transfer of membrane proteins between the cells would require dedicated study. We also challenged our two-color cell tracking approach in video imaging for 1h. We could clearly see fast dynamics of cell membranes of both green (MemGraft-Cy3) and red (MemGraft-Cy5) cell populations (Video S2). Although MemGraft-Cy5 showed some photobleaching in contrast to MemGraft-Cy3, both probes allowed tracking of cell migration without any visual impact on their behavior. The latter confirmed negligible phototoxicity of these probes, which is essential for the multi-color long-term cell tracking.

**Figure 8.**
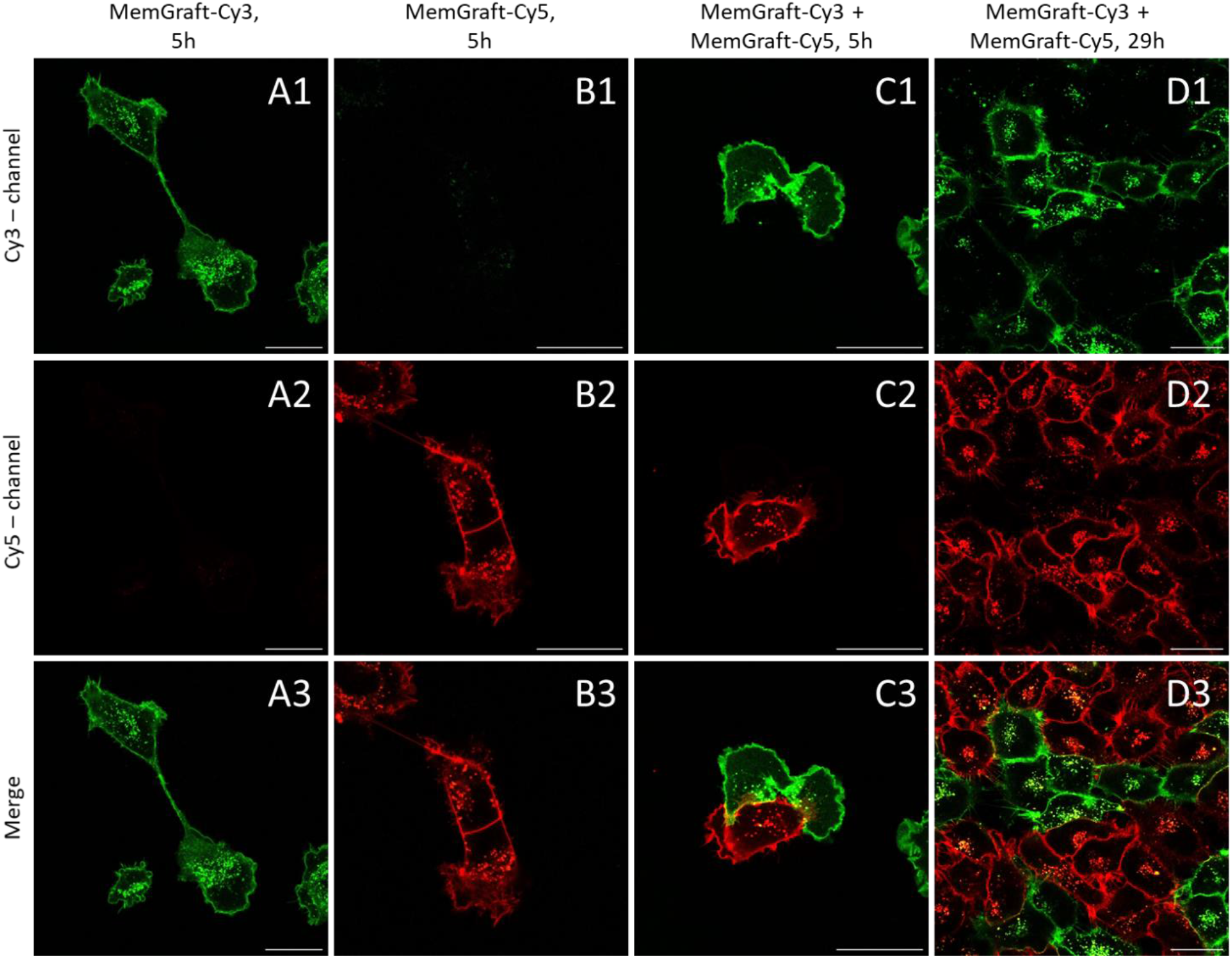
Co-seeding of cells stained with MemGraft-Cy3 and MemGraft-Cy5 (C): after 5h ; (D): after 29h. Cells stained only with MemGraft-Cy3 (500 nM) or MemGraft-Cy5 (500 nM) are shown respectively in A and B Scale bar: 30 μm.

### Cell barcoding

Inspired by the capacity of MemGraft probes to track cells in two colors without exchange of the Cy3- and Cy5-analogues, we explored the possibility to barcode the cells in multiple colors using these probes. The covalent labelling of PM could in principle enable co-staining of PM with different combinations of green (MemGraft-Cy3) and red (MemGraft-Cy5) probes, thus generating a characteristic barcode for each cell population. To this end, we labelled cells using five probe combinations of MemGraft-Cy3 / MemGraft-Cy5 molar %: (1) 100%/0%, (2) 75%/25%, (3) 50%/50%, (4) 25%/75% and (5) 0%/100%. First, individual cell populations labelled with these combinations were imaged using green and red channels, corresponding to Cy3 and Cy5 dyes. Once the images from individual channels were built, the merged images were constructed by simple combination of the two channels (Figure S12). As a result, we observed that each population was characterized by its own color code, which was homogenous within the cell population: green, lime, yellow, orange and red (Figure 9A). This result indicates that the two probes stain cells in a highly homogenous manner and with a similar efficiency. Then, all five cell populations were detached and then co-incubated together for 3 h and imaged by the same method (Figure 9B). We found that in the cells presenting different color barcodes were simultaneously present on the surface (Figure 9C). Importantly, all five populations with green, lime, yellow, orange and red barcodes could be readily identified on these images. These observations indicate that after cell detachment of co-culture, the barcode was preserved within the individual cells and it did not exchange within different cells. Thus, MemGraft dyes enable barcoding of cells at the level of PM and further tracking of multiple (at least 5) cell populations simultaneously, using just two probes.

**Figure 9.**
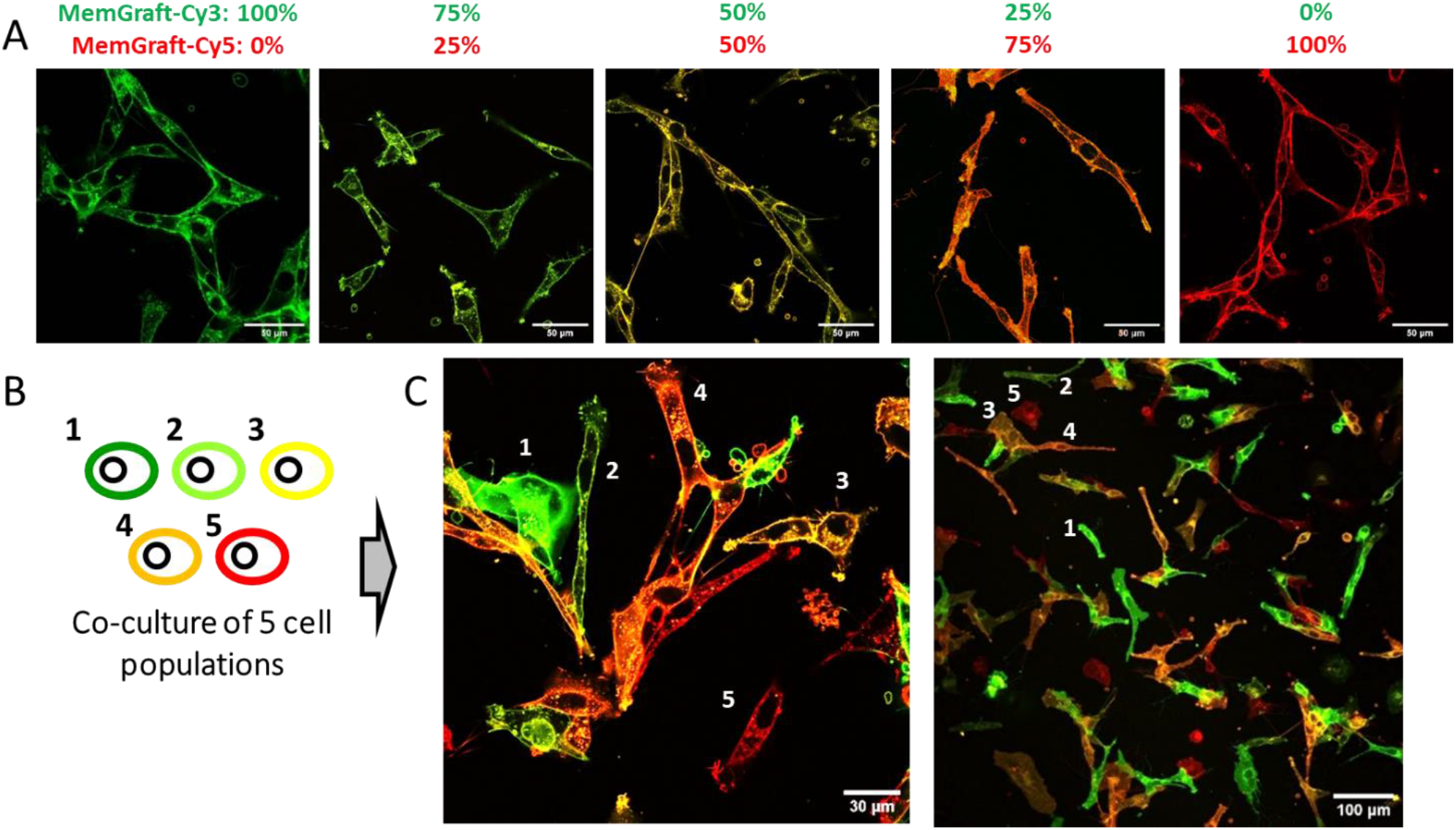
(A) Confocal fluorescence microscopy of U87 cells stained with growing ratio of concentration MemGraft-Cy3 / MemGraft-Cy5 ranging 100, 75, 50, 25, 0 (left to right: green, lime, yellow, orange, red)). Total dye concentration was 1 μm in each case. Scale bars: 100 μm. (B) Scheme showing that five cell populations where mixed together and co-cultured. (C) Confocal fluorescence images of U87 cells of five barcoded cell populations mixed together and incubated for 3 h before imaging. Total dye concentration was 1 μm in each case. Scale bars: 30 and 100 μm. The numbers indicate the cells bearing one of the five barcodes.

### Cell biotinylation and manipulation

The capacity of MemGraft probes to label covalently live cells at their plasma membrane opens unique opportunities in cell surface engineering, which is a rapidly developing research direction.^60,61^ To address this point, we designed MemGraft-Cy3 derivative bearing biotin residue through a PEG3 linker (MemGraft-Cy3-biotin, Figure 10A). Biotin is an attractive plug-and-play unit, which enables specific conjugation and manipulation using biotin-streptavidin coupling. Incubation of cells with MemGraft-Cy3-biotin resulted in specific plasma membrane labeling, which showed that modification of MemGraft-Cy3 with PEG3 and biotin units did not alter its PM-specific labelling (Figure 10B). Next, we verified whether the attached biotin unit is capable to interact with streptavidin. To this end, cells labelled with MemGraft-Cy3-biotin were incubated with strepatividin-Cy5 conjugate and then imaged at three channels: (1) the green (Cy3) channel to visualize MemGraft-Cy3 part; (2) red (Cy5) emission channel with Cy3 excitation in order to eventually observe Forster Resonance Energy Transfer (FRET) from MemGraft-Cy3 to attached strepatvidin-Cy5 (FRET channel) and (3) at the red channel with Cy5 excitation in order to detect streptavidin-Cy5 part (direct Cy5 channel). We found the strong membrane-specific signal at the Cy5 channel with Cy3-exciation and its colocalization well the with the signal at the green (Cy3) channel (Figure 10C). Direct excitation of Cy5 also produced membrane-specific signal, but the background noise was much more significant. In case of control dye MemGraft-Cy3 (without biotin) a much weaker signal was observed at the FRET channel, whereas in the direct Cy5 channel only background noise was observed. Another control experiment with non-labelled cells treated with strepatvidin-Cy5 showed no signal from cell membranes in any of the studied channels (Figure S13). These observations provide a strong evidence of specific attachment of streptavidin-Cy5 to the cells labelled with MemGraft-Cy3-biotin. The latter leads to the close proximity of Cy5 label with the MemGraft-Cy3 part, leading to observed FRET signal. In contrast, a much weaker signal observed for the FRET signal with control MemGraft-Cy3 corresponds to the leakage of the Cy3 emission into the Cy5 emission channel. It should be noted that direct Cy5 channel also confirmed the attachment of streptavidin-Cy5, but the quality of the signal was much worse because in this case we excite all streptaividin-Cy5 conjugates, including those bound non-specifically to the glass surface. Thus, FRET approach provides more specific signal of streptavidin-Cy5 fraction bound to the plasma membrane.

**Figure 10.**
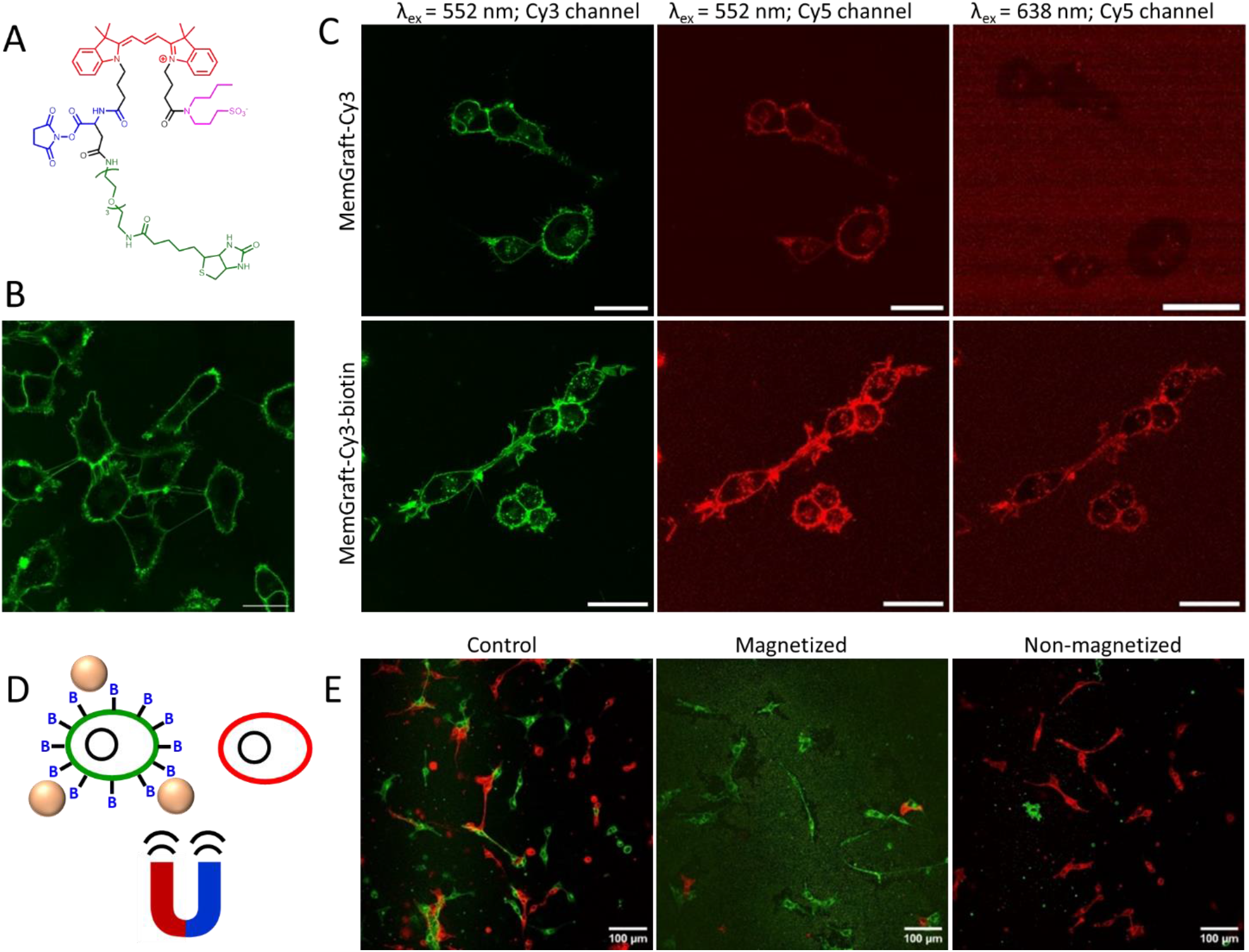
(A) Chemical structure of MemGraft-Biotin-Cy3. (B) Confocal fluorescence microscopy of U87 cells stained with MemGraft-Biotin-Cy3. Dye concentration was 500 nM. Scale bars: 30 μm. (C) Confocal fluorescence microscopy of U87 cells stained with MemGraft-Cy3 (upper panels) and MemGraft-Cy3-biotin (lower panels) and incubated with streptavidin-Cy5 conjugate. Left panel: the image is recorded with excitation of Cy3 (552 nm) and emission of Cy3. Middle panel: the image is recorded with excitation of Cy3 (552 nm) and emission of Cy5. Right panel: the image is recorded with excitation of Cy5 (638 nm) and emission of Cy5. MemGraft probes concentration was 500 nM in each case. The cells were washed and incubated for 30 min with streptavidin-Cy5 (0.1 mg/mL). Scale bars: 30 μm. (D) Scheme of magnetic separation of biotinylated cells (in green) using magnetic beads (colored in beige) from non-biotinylated ones (in red). (E) Confocal fluorescence microscopy of U87 cells stained separately with MemGraft-Biotin-Cy3 and MemGraft-Cy5 and mixed together. Dye concentration was 500 nM. Left panel: control cells mixed without further treatment. Middle panel: the magnetized fraction of cells incubated with streptavidin magnetic beads and isolated using a magnetic stand. Right panel: non-magnetized cells from the supernatant after incubation with streptavidin magnetic beads and placing on a magnetic stand. Scale bars: 100 μm.

Ultimately, we tested whether MemGraft-based cell-biotinylation approach could be used for cell manipulation. To this end, we applied streptavidin coated magnetic beads, which are commonly used to separate biotinylated molecules or cells from non-biotinylated ones. In our experimental setup, one cell population was labelled with MemGraft-Cy3-biotin, while the other one with non-biotinylated MemGraft-Cy5 (Figure 10D). Then, the cells were detached and mixed together. This control conditions gave two cell populations, corresponding to biotinylated cells stained in green by MemGraft-Cy3-biotin and non-biotinylated ones stained in red by MemGraft-Cy5, as evidenced by two-color fluorescence imaging (Figure 10E). In parallel, the suspension of the two cell populations was exposed to a magnetic field, which resulted in rapid precipitation of some fraction of cells, which was isolated from supernatant (Figure 10D). Then, the isolated magnetized and remaining (non-magnetized) supernatant suspensions were deposited on glass slides and imaged. Remarkably, the magnetized fraction showed predominantly green emission of the cells, whereas the remaining non-magnetized suspension was stained in red (Figure 10E). These experiments provided the proof of concept of cell manipulation coupled to fluorescence imaging using MemGraft approach in combination with biotin-streptavidin coupling strategy. The advantage of using MemGraft strategy is linked to the higher labelling density, as it was evidenced from much more efficient cell labelling by MemGraft compared to the classical NHS labelling approach (see Figures 3 and 4). Moreover, the fluorescence modality of MemGraft allows direct observation of the cell surface modification and its possible impact on the plasma membrane morphology by fluorescence microscopy.

## Conclusion

Permanent labelling of cell plasma membranes remains a challenge because of the non-covalent nature of current fluorescent membrane probes. Here, we report a concept of lipid-directed covalent labelling of plasma membranes. We propose to exploit transient binding to biomembrane surface as a driving force creating a high local concentration of the dye in proximity of membrane proteins, which can catalyze ligation to membrane proteins. We designed a new family of dyes, named MemGraft based on a cyanine dye that bears a low-affinity membrane anchor and an activated ester (or maleimide) on two ends of the dye. We found that the low-affinity anchor plays a crucial role in ensuring efficient PM labelling, as a control compound without this group failed to stain PM. Moreover, MemGraft probes with activated ester and maleimide were found superior to corresponding commercial NHS-ester and maleimide Cy3-based labels. These data confirmed that the anchor ensures transient binding to the plasma membrane that accelerates labelling of amino or thiol groups of membrane proteins in the proximity of lipid membranes. The reactivity of the NHS-ester was found crucial, as less reactive fluorophenyl activated esters showed strong internalization and inefficient PM labelling. SDS-page confirmed that multiple membrane proteins were labeled with MemGraft-Cy3 probes (with activated ester and maleimide), thus confirming that our strategy of lipid-driven covalent labelling of the cell surface operated through the non-selective conjugation to the membrane proteins. In comparison to conventional non-covalent staining of PM by reference probes (e.g. MemBright), the MemGraft approach revealed several critical advantages: resistance to full culture medium with serum in long-term cellular experiments and compatibility with fixation and permeabilization. Additionally, these probes enabled long-term labelling of cells and co-culture of two cell populations stained in different colors without dye exchange, which was not possible before with conventional membrane probes. The latter allowed imaging of cell-cell interactions and visualization of cell-cell contacts. Moreover, combination of different ratios of MemGraft-Cy3 and MemGraft-Cy5 probes (NHS-esters) enabled long term cell barcoding in at least 5 color codes, which opens new opportunities in tracking and visualizing multiple cells populations. This approach is analogous to a well-established “Brainbow” method,^62,63^ but it does not require genetic modification of the cells. Ultimately, based on MemGraft strategy, we designed a fluorescent probe for efficient biotinylation of cell surface, which enabled cell manipulation using streptavidin-coated magnetic beads. The developed MemGraft concept opens a new chapter in the design of advanced PM probes and, in a more general sense, proposes an efficient approach for the covalent functionalization of any biomembrane surface and cell surface engineering.

## Supporting Information

Supporting information is available. The authors have cited the following references within the Supporting Information.^11,38^

## Acknowledgments

This work was supported by the Interdisciplinary Thematic Institute SysChem, via the IdEx Unistra (ANR-10-IDEX-0002), the CSC Graduate School (CSC-IGS ANR-17-EURE-0016) within the French Investments for the Future Program and Agence Nationale de la Recherche (ANR) AmpliSens ANR-21-CE42-0019-01. We thank Dr. Denis Dujardin for help with video microscopy experiments.

## Author contributions

NA performed majority of the experiments, analyzed the results and contributed to manuscript writing. RP contributed to the probe design, SDS-Page and flow cytometry experiments. ASK designed and supervised the project, analyzed the results and wrote the manuscript.

## Conflict of interest

The authors declare no conflict of interest.

## Graphical Abstract

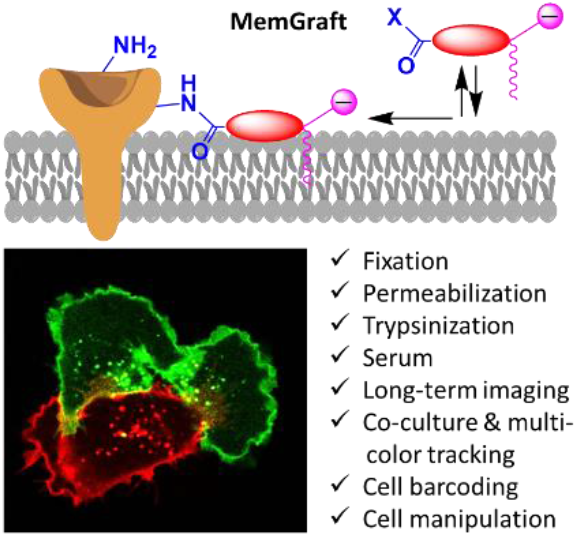

## Notes

### Competing Interest Statement

The authors have declared no competing interest.

