## Supplemental information for "Lipid-directed covalent fluorescent labeling of plasma membranes for long-term imaging, barcoding and manipulation of cells"

### Materials and methods

#### General Methods and Materials

All the reagents were purchased from Sigma-Aldrich, Alfa Aesar, or TCI and used without any purification. Cy3 and Cy5 diacids, 3-(butylammonio)propane-1-sulfonate were synthesized as described previously.[1,2] Disulfo-Cy3 NHS ester was purchased from Tebu-Bio. Disulfo-Cy3-maleimide was purchased from Interchim. MilliQ-water (Millipore) was used in all experiments. NMR spectra were recorded on a BrukerAvance III 400 MHz spectrometer and a BrukerAvance III 500 MHz spectrometer. Mass spectra were obtained using an Agilent Q-TOF 6520 mass spectrometer with electrospray ionization and a Bruker Microflex LRF system for MALDI-TOF MS. Absorption and emission spectra were recorded on an Agilent Cary 5000 UV−vis−NIR spectrophotometer and an Edinburgh FS5 spectrofluorometer, correspondingly.

### Synthesis

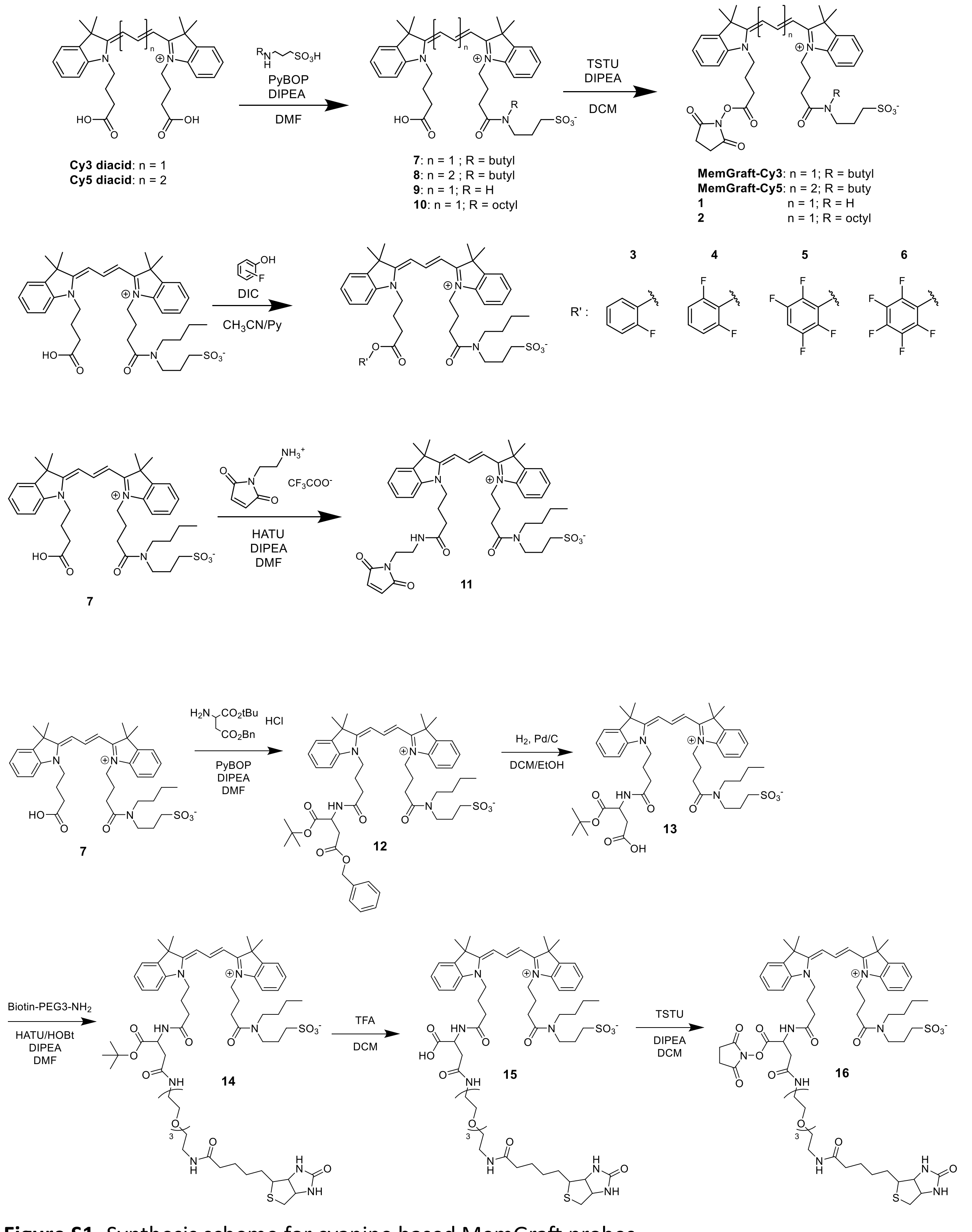


**Figure S1.** Synthesis scheme for cyanine based MemGraft probes.

**3-(N-butyl-4-(2-((E)-3-((Z)-1-(3-carboxypropyl)-3,3-dimethylindolin-2-ylidene)prop-1-en-1-yl)-3,3-dimethyl-3H-indol-1-ium-1-yl)butanamido)propane-1-sulfonate (7).**

The synthesis is adapted from the protocol decreased elsewhere.[1] Compound Cy3 diacid chloride (110 mg, 204.8 µmol) was dissolved in dry DMF together with 3-(butylammonio)propane-1-sulfonate (40.0 mg, 204.8 µmol) and DIPEA (125 µL, 716.9 µmol). The mixture was cooled to 0°C and PyBOP (106.6 mg, 204.8 µmol) was added. The mixture was allowed to stir at room temperature for 4h (controlled by TLC). The reaction was quenched with water and the mixture was evaporated *in vacuo*. The crude product was purified by flash chromatography on silica $SiO_2$ gel with gradual eluting with DCM/methanol from 98:2 to 85:15 v/v %. Yield: 50 mg (36%) as a dark magenta solid. $^1$H NMR (500 MHz, MeOD) δ ppm: 8.43 (td, *J* = 13.4, 6.6 Hz, 1H), 7.46 – 7.41 (m, 2H), 7.38 – 7.30 (m, 4H), 7.20 (tt, *J* = 7.6, 2.6 Hz, 2H), 6.55 – 6.41 (m, 2H), 4.12 (td, *J* = 10.5, 6.5 Hz, 4H), 3.45 (tq, *J* = 12.0, 5.4 Hz, 2H), 3.31 – 3.18 (m, 4H), 2.51 – 2.43 (m, 2H), 2.00 (dd, *J* = 41.2, 22.8, 9.7, 6.7 Hz, 7H), 1.66 (dd, *J* = 3.9, 1.4 Hz, 12H), 1.42 (tt, *J* = 8.8, 6.4 Hz, 2H), 1.23 – 1.12 (m, 4H), 0.82 (dd, *J* = 15.2, 7.6 Hz, 3H). $^{13}$C NMR (126 MHz, MeOD) δ ppm: 174.72, 172.43, 150.95, 150.77, 141.96, 140.82, 128.66, 128.27, 125.37, 125.03, 122.14, 119.07, 111.16, 109.00, 102.68, 49.26, 48.53, 46.55, 45.65, 44.68, 43.31, 42.44, 30.97, 30.36, 29.34, 28.67, 28.24, 26.98, 24.22, 23.91, 23.01, 22.52, 22.23, 12.87. HRMS (ESI), m/z: [M+H]+ calcd for $C_{38}H_{52}N_3O_6S^+$, 678.3571; found, 678.3585.

**3-(N-butyl-4-(2-((1E,3E)-5-((Z)-1-(3-carboxypropyl)-3,3-dimethylindolin-2-ylidene)penta-1,3-dien-1-yl)-3,3-dimethyl-3H-indol-1-ium-1-yl)butanamido)propane-1-sulfonate (8).**

The synthesis is adapted from the protocol decreased elsewhere.[1] Compound Cy5 diacid chloride (98 mg, 174.0 µmol) was dissolved in dry DMF together with 3-(butylammonio)propane-1-sulfonate (34.5 mg, 174.0 µmol) and DIPEA (106 µL, 609.1 µmol). The mixture was cooled to 0°C and PyBOP (90.6 mg, 174.0 µmol) was added. The mixture was allowed to stir at room temperature for 4h (controlled by TLC). The reaction was quenched with water and the mixture was evaporated *in vacuo*. The crude product was purified by flash chromatography on silica $SiO_2$ gel with gradual eluting with DCM/methanol from 98:2 to 85:15 v/v %. Yield: 17.5 mg (14%) as a dark blue solid. $^1$H NMR (400 MHz, MeOD) δ ppm: 8.12 (t, *J* = 12.9 Hz, 2H), 7.37 (dt, *J* = 7.2, 2.6 Hz, 2H), 7.33 – 7.20 (m, 4H), 7.13 (dtt, *J* = 9.1, 3.5, 1.3 Hz, 2H), 6.58 (dt, *J* = 23.6, 12.3 Hz, 1H), 6.46 – 6.17 (m, 2H), 4.08 (q, *J* = 10.8 Hz, 4H), 3.45 – 3.38 (m, 2H), 2.79 – 2.72 (m, 2H), 2.41 (dt, *J* = 21.7, 6.4 Hz, 3H), 1.97 (q, *J* = 8.1 Hz, 6H), 1.60 (d, *J* = 2.3 Hz, 12H), 1.41 (dd, *J* = 10.0, 5.4 Hz, 2H), 1.29 – 1.12 (m, 6H), 0.84 – 0.78 (m, 3H). $^{13}$C NMR (101 MHz, MeOD) δ ppm: 172.06, 171.69, 170.93, 170.80, 152.80, 152.48, 140.63, 139.85, 139.75, 139.68, 126.90, 124.19, 123.51, 123.31, 120.50, 109.26, 109.11, 101.51, 52.97, 47.66, 47.24, 46.96, 45.00, 44.11, 41.43, 40.92, 29.03, 27.74, 27.03, 26.71, 25.05, 22.68, 21.72, 21.10, 20.90, 20.67, 18.36, 18.23, 11.36, 10.27. HRMS (ESI), m/z: [M+H]+ calcd for $C_{40}H_{53}N_3O_6S$, 703.3655; found, 703.3642.

**3-(4-(2-((E)-3-((Z)-1-(4-((2,5-dioxopyrrolidin-1-yl)oxy)-4-oxobutyl)-3,3-dimethylindolin-2-ylidene)prop-1-en-1-yl)-3,3-dimethyl-3H-indol-1-ium-1-yl)butanamido)propane-1-sulfonate (9).**

Compound Cy3 diacid (40 mg, 68.7 µmol) was dissolved in dry DMF together with 3-aminopropane-1-sulfonic acid (10.5 mg, 76.0 µmol) and DIPEA (43 µL, 246.8 µmol). The mixture was cooled to 0°C and PyBOP (40 mg, 76.0 µmol) was added. The mixture was allowed to stir at room temperature for 4h (controlled by TLC). The reaction was quenched with water and the mixture was evaporated *in vacuo*. The crude product was purified by preparative TLC ($SiO_2$, DCM/MeOH, 85:15 v/v %). Yield: 15 mg (35%) as a dark magenta solid. $^1H$ NMR (400 MHz, MeOD) δ ppm: 8.61 – 8.51 (m, 1H), 7.61 – 7.52 (m, 2H), 7.50 – 7.38 (m, 4H), 7.38 – 7.26 (m, 2H), 6.71 – 6.55 (m, 2H), 4.22 (p, *J* = 5.9 Hz, 4H), 3.35 (s, 4H), 2.90 (dt, *J* = 29.0, 7.3 Hz, 2H), 2.64 – 2.47 (m, 4H), 2.13 (pd, *J* = 9.7, 5.0 Hz, 6H), 2.03 – 1.84 (m, 2H), 1.77 (d, *J* = 1.7 Hz, 12H). $^{13}C$ NMR (101 MHz, MeOD) δ 174.72, 173.46, 173.00, 150.94, 141.93, 140.82, 128.63, 128.10, 125.38, 122.12, 111.10, 102.56, 54.68, 50.90, 49.27, 48.94, 43.30, 43.07, 38.10, 31.75, 29.96, 28.17, 26.93, 24.81, 22.75, 22.00, 13.03. HRMS (ESI), m/z: [M+H]+ calcd for $C_{34}H_{43}N_3O_6S$, 621.2873; found, 621.2879.

**3-(4-(2-((E)-3-((Z)-1-(3-carboxypropyl)-3,3-dimethylindolin-2-ylidene)prop-1-en-1-yl)-3,3-dimethyl-3H-indol-1-ium-1-yl)-N-octylbutanamido)propane-1-sulfonate (10)**

Compound Cy3 diacid chloride (110 mg, 204.8 µmol) was dissolved in dry DMF together with 3-(octylammonio)propane-1-sulfonate (40.0 mg, 204.8 µmol) and DIPEA (125 µL, 716.9 µmol). The mixture was cooled to 0°C and PyBOP (106.6 mg, 204.8 µmol) was added. The mixture was allowed to stir at room temperature for 4h (controlled by TLC). The reaction was quenched with water and the mixture was evaporated *in vacuo*. The crude product was purified by flash chromatography on silica $SiO_2$ gel with gradual eluting with DCM/methanol from 98:2 to 85:15 v/v %. Yield: 51 mg (36%) as a dark magenta solid. $^1H$ NMR (400 MHz, $CDCl_3$) δ 8.45 – 8.31 (m, 1H), 7.39 (q, *J* = 6.8 Hz, 3H), 7.32 (dd, *J* = 7.4, 2.5 Hz, 3H), 7.22 (tdd, *J* = 8.8, 6.4, 3.9 Hz, 2H), 6.97 – 6.77 (m, 2H), 4.22 (s, 2H), 3.64 – 3.49 (m, 2H), 3.29 (d, *J* = 7.6 Hz, 2H), 3.19 (s, 4H), 3.16 – 3.05 (m, 4H), 2.93 (d, *J* = 8.7 Hz, 4H), 2.81 (t, *J* = 7.0 Hz, 2H), 2.76 – 2.62 (m, 2H), 2.42 – 2.31 (m, 4H), 2.18 – 2.00 (m, 6H), 1.69 (s, 12H), 1.35 – 1.10 (m, 10H), 0.82 (s, 3H). $^{13}C$ NMR (101 MHz, $CDCl_3$) δ 173.67, 172.01, 150.79, 141.93, 141.90, 140.43, 140.40, 129.19, 129.13, 125.43, 125.30, 121.92, 121.81, 111.84, 111.38, 103.99, 54.11, 49.13, 48.97, 48.78, 48.17, 47.41, 47.22, 46.69, 45.92, 44.20, 42.35, 31.80, 31.72, 30.10, 29.69, 29.46, 29.42, 29.27, 29.23, 29.16, 29.10, 28.25, 28.16, 28.12, 27.93, 27.08, 26.93, 26.74, 25.97, 24.92, 23.54, 22.62, 22.59, 21.85, 19.03, 14.07, 14.06. HRMS (ESI), m/z: [M+H]+ calcd for $C_{42}H_{59}N_3O_6S$, 733.4125; found, 733.4134.

**3-(N-butyl-4-(2-((E)-3-((Z)-1-(4-(2-fluorophenoxy)-4-oxobutyl)-3,3-dimethylindolin-2-ylidene)prop-1-en-1-yl)-3,3-dimethyl-3H-indol-1-ium-1-yl)butanamido)propane-1-sulfonate (3).**

Compound **7** (20 mg, 29.5 µmol) was added to a stirred solution of DIC (5 µL, 35.4 µmol) in $CH_3CN$/Pyridine (9:1). The mixture was allowed to stir for 15 minutes. 2-fluorophenol (3.5 µL, 35.4 µmol) was slowly added and the reaction was stirred for 12h. The solvent was evaporated *in vacuo* and the crude product was purified by flash chromatography on silica $SiO_2$ gel with gradual eluting with DCM/methanol from 98:2 to 90:10 v/v %. Yield: 9 mg (40%) as a dark magenta solid. $^1H$ NMR (500 MHz, $CDCl_3$) δ ppm: 8.37 (t, *J* = 13.4 Hz, 1H), 7.33 – 7.24 (m, 6H), 7.18 – 7.06 (m, 8H), 4.44 – 4.37 (m, 2H), 4.16 (t, *J* = 7.6 Hz, 2H), 3.69 (dd, *J* = 9.9, 6.6 Hz, 2H), 3.26 (t, *J* = 7.5 Hz, 2H), 3.12 (t, *J* = 6.4 Hz, 2H), 2.94 – 2.89 (m, 4H), 2.23 – 2.11 (m, 4H), 1.65 (d, *J* = 4.5 Hz, 12H), 1.42 (ddt, *J* = 9.2, 7.7, 3.5 Hz, 2H), 1.27 – 1.16 (m, 4H), 0.83 (t, *J* = 7.3 Hz, 3H). $^{13}C$ NMR (126 MHz, $CDCl_3$) δ ppm: 174.00,

173.85, 172.52, 171.68, 151.48, 142.16, 141.91, 140.59, 140.45, 138.09, 129.09, 128.94, 127.03, 125.19, 124.51, 123.92, 121.86, 116.68, 116.53, 111.24, 105.10, 104.53, 48.96, 47.79, 46.91, 44.34, 43.50, 36.46, 33.95, 31.42, 31.03, 30.10, 28.19, 25.81, 25.66, 24.98, 24.63, 20.18, 13.88. HRMS (ESI), m/z: [M+H]+ calcd for $C_{44}H_{54}FN_3O_6S$, 771.3717; found, 771.3706.

**3-(N-butyl-4-(2-((E)-3-((Z)-1-(4-(2,6-difluorophenoxy)-4-oxobutyl)-3,3-dimethylindolin-2-ylidene)prop-1-en-1-yl)-3,3-dimethyl-3H-indol-1-ium-1-yl)butanamido)propane-1-sulfonate (4).**

Compound **7** (21 mg, 30.9 µmol) was added to a stirred solution of DIC (6 µL, 37 µmol) in $CH_3CN$/Pyridine (9:1). The mixture was allowed to stir for 15 minutes. 2,4-difluorophenol (3.5 µL, 37 µmol) was slowly added and the reaction was stirred for 12h. The solvent was evaporated *in vacuo* and the crude product was purified by preparative TLC ($SiO_2$, DCM/MeOH, 85:15 v/v%). Yield: 20 mg (82%) as a dark magenta solid. $^1$H NMR (400 MHz, $CDCl_3$) δ ppm: 8.43 (t, *J* = 13.4 Hz, 1H), 7.44 – 7.27 (m, 6H), 7.25 – 7.08 (m, 5H), 6.96 – 6.80 (m, 2H), 4.50 – 4.41 (m, 2H), 4.22 (t, *J* = 7.6 Hz, 2H), 3.76 (dd, *J* = 10.0, 6.4 Hz, 2H), 3.32 (t, *J* = 7.5 Hz, 2H), 3.20 (t, *J* = 6.4 Hz, 2H), 3.02 – 2.95 (m, 2H), 2.85 (t, *J* = 8.0 Hz, 2H), 2.21 (dd, *J* = 23.6, 6.7 Hz, 4H), 1.71 (d, *J* = 3.6 Hz, 12H), 1.49 (tt, *J* = 8.0, 6.4 Hz, 3H), 1.29 – 1.24 (m, 4H), 0.89 (t, *J* = 7.3 Hz, 3H). $^{13}$C NMR (101 MHz, $CDCl_3$) δ ppm: 173.93, 172.56, 171.78, 151.51, 142.16, 141.91, 140.60, 140.46, 128.96, 125.21, 124.47, 121.88, 105.26, 105.05, 104.77, 104.58, 48.95, 30.16, 29.97, 29.70, 28.21, 25.82, 24.68, 22.30, 20.19, 13.89. HRMS (ESI), m/z: [M+H]+ calcd for $C_{44}H_{53}F_2N_3O_6S$, 789.3623; found, 789.3644.

**3-(N-butyl-4-(2-((E)-3-((Z)-3,3-dimethyl-1-(4-oxo-4-(2,3,5,6-tetrafluorophenoxy)butyl)indolin-2-ylidene)prop-1-en-1-yl)-3,3-dimethyl-3H-indol-1-ium-1-yl)butanamido)propane-1-sulfonate (5)**

Compound **7** (24 mg, 35.4 µmol) was added to a stirred solution of DIC (6.6 µL, 43 µmol) in $CH_3CN$/Pyridine (9:1). The mixture was allowed to stir for 15 minutes. 2,3,5,6-tetrafluorophenol (7 mg, 43 µmol) was slowly added and the reaction was stirred for 12h. The solvent was evaporated *in vacuo* and the crude product was purified by preparative TLC ($SiO_2$, DCM/MeOH, 85:15 v/v%). Yield: 15 mg (52%) as a dark magenta solid. $^1$H NMR (400 MHz, $CDCl_3$) δ ppm: 8.48 – 8.34 (m, 1H), 7.49 – 7.26 (m, 6H), 7.24 – 7.07 (m, 3H), 6.98 (tt, *J* = 9.9, 7.1 Hz, 1H), 6.44 (tt, *J* = 10.2, 6.9 Hz, 1H), 4.47 – 4.38 (m, 2H), 4.21 (t, *J* = 7.7 Hz, 2H), 3.84 (p, *J* = 6.5 Hz, 2H), 3.75 (q, *J* = 7.4 Hz, 2H), 3.34 (q, *J* = 7.3 Hz, 2H), 3.27 (t, *J* = 6.6 Hz, 2H), 3.04 – 2.94 (m, 2H), 2.89 – 2.78 (m, 2H), 2.33 – 2.23 (m, 2H), 2.23 – 2.08 (m, 4H), 1.71 (d, *J* = 2.3 Hz, 12H), 1.49 (p, *J* = 7.7 Hz, 2H), 1.36 – 1.17 (m, 6H), 0.90 (q, *J* = 7.1 Hz, 3H). $^{13}$C NMR (101 MHz, $CDCl_3$) δ ppm: 174.08, 173.95, 172.78, 170.18, 157.22, 151.54, 142.06, 141.80, 140.56, 129.04, 125.29, 121.92, 104.85, 104.52, 104.21, 103.11, 102.88, 94.94, 94.71, 94.48, 53.43, 49.24, 48.98, 47.75, 46.94, 45.81, 44.32, 43.29, 42.18, 30.94, 30.10, 29.79, 25.66, 24.60, 23.46, 22.22, 20.19, 14.11, 13.87. HRMS (ESI), m/z: [M+H]+ calcd for $C_{44}H_{51}F_4N_3O_6S$, 825.3435; found, 825.3438.

**3-(N-butyl-4-(2-((E)-3-((Z)-3,3-dimethyl-1-(4-oxo-4-(perfluorophenoxy)butyl)indolin-2-ylidene)prop-1-en-1-yl)-3,3-dimethyl-3H-indol-1-ium-1-yl)butanamido)propane-1-sulfonate (6).**

Compound **7** (22 mg, 32.5 µmol) was added to a stirred solution of DIC (6 µL, 39 µmol) in $CH_3CN$/Pyridine (9:1). The mixture was allowed to stir for 15 minutes. 2,3,4,5,6-pentafluorophenol

(6.5 mg, 39 µmol) was slowly added and the reaction was stirred for 12h. The solvent was evaporated *in vacuo* and the crude product was purified by preparative TLC ($SiO_2$, DCM/MeOH, 85:15 v/v %). Yield: 18 mg (67%) as a dark magenta solid. $^1H$ NMR (400 MHz, $CDCl_3$) δ ppm: 8.49 – 8.33 (m, 1H), 7.35 (tdd, *J* = 14.8, 12.6, 7.3 Hz, 5H), 7.25 – 6.94 (m, 4H), 6.77 (dd, *J* = 30.8, 13.4 Hz, 1H), 4.30 (s, 2H), 4.20 (t, *J* = 7.8 Hz, 2H), 3.84 (p, *J* = 6.4 Hz, 3H), 3.77 – 3.67 (m, 2H), 3.34 (q, *J* = 7.7 Hz, 2H), 2.99 (dd, *J* = 7.5, 3.7 Hz, 2H), 2.82 (dd, *J* = 14.6, 8.3 Hz, 3H), 2.69 (t, *J* = 6.9 Hz, 1H), 2.20 – 2.12 (m, 4H), 1.71 (t, *J* = 1.6 Hz, 12H), 1.30 – 1.23 (m, 4H), 0.90 (td, *J* = 7.4, 5.6 Hz, 3H). $^{13}C$ NMR (101 MHz, $CDCl_3$) δ ppm: 174.32, 174.15, 173.98, 173.75, 172.87, 170.17, 151.51, 141.99, 141.72, 140.55, 129.07, 125.66, 121.84, 111.71, 111.26, 111.14, 104.67, 104.36, 103.84, 49.27, 47.71, 46.94, 46.69, 44.26, 43.22, 42.25, 30.86, 29.69, 28.14, 25.57, 24.58, 23.89, 23.44, 13.86. HRMS (ESI), m/z: [M+H]+ calcd for $C_{44}H_{50}F_5N_3O_6S$, 843.334; found, 843.3339.

**3-(N-butyl-4-(2-((E)-3-((Z)-1-(4-((2-(2,5-dioxo-2,5-dihydro-1H-pyrrol-1-yl)ethyl)amino)-4-oxobutyl)-3,3-dimethylindolin-2-ylidene)prop-1-en-1-yl)-3,3-dimethyl-3H-indol-1-ium-1-yl)butanamido)propane-1-sulfonate (11)**

Compound **7** (25 mg, 36.9 µmol) was dissolved in DMF together with N-(2-aminoethyl)maleimide trifluoroacetate salt (10.5 mg, 42 µmol) and DIPEA (23 µL, 132 µmol). The mixture was cooled to 0°C and HATU (16 mg, 42 µmol) was added. The mixture was allowed to stir at room temperature for 4h (controlled by TLC). The reaction was quenched with water and the mixture was evaporated *in vacuo*. The crude product was purified by flash chromatography on silica $SiO_2$ gel with gradual eluting with DCM/methanol from 98:2 to 85:15 v:v%. Yield: 24 mg (81%) as a dark red solid. $^1H$ NMR (400 MHz, $CDCl_3$) δ 8.39 – 8.26 (m, 2H), 7.24 (m, 7H), 6.94 (d, *J* = 13.4 Hz, 1H), 6.80 (d, *J* = 13.3 Hz, 1H), 6.60 (s, 2H), 4.13 (dt, *J* = 14.5, 7.4 Hz, 4H), 3.63 (t, *J* = 5.7 Hz, 4H), 3.39 (t, *J* = 5.7 Hz, 2H), 3.26 (t, *J* = 7.6 Hz, 2H), 2.85 (t, *J* = 5.6 Hz, 2H), 2.76 (t, *J* = 7.7 Hz, 2H), 2.57 (q, *J* = 5.2 Hz, 2H), 1.64 (s, *J* = 2.6 Hz, 12H), 1.43 (p, *J* = 7.8 Hz, 2H), 1.29 – 1.14 (m, 6H), 0.84 (t, *J* = 7.3 Hz, 3H). $^{13}C$ NMR (126 MHz, $CDCl_3$) δ 174.15, 173.67, 170.91 (d, *J* = 2.7 Hz), 151.15, 150.36, 141.82, 140.49 (d, *J* = 3.1 Hz), 140.34, 134.11, 129.10 (d, *J* = 15.0 Hz), 125.49, 125.26, 121.86 (d, *J* = 10.7 Hz), 111.79, 111.31, 104.62, 103.85, 53.43, 49.04 (d, *J* = 11.4 Hz), 47.75, 46.95, 44.03 (d, *J* = 12.9 Hz), 38.22, 38.05, 37.60 (d, *J* = 10.7 Hz), 33.12, 32.91, 31.92, 30.92, 30.51 (d, *J* = 13.0 Hz), 30.05 (d, *J* = 3.0 Hz), 29.69, 29.27, 28.21, 28.10, 25.30, 24.30 (d, *J* = 21.0 Hz), 20.19, 13.87.

HRMS (ESI), m/z: [M+H]+ calcd for $C_{44}H_{57}N_5O_7S$, 799.3979; found, 799.3986.

**3-(4-(2-((E)-3-((Z)-1-(4-((4-(benzyloxy)-1-(tert-butoxy)-1,4-dioxobutan-2-yl)amino)-4-oxobutyl)-3,3-dimethylindolin-2-ylidene)prop-1-en-1-yl)-3,3-dimethyl-3H-indol-1-ium-1-yl)-N-butylbutanamido)propane-1-sulfonate (12)**

Compound **7** (80 mg, 118 µmol) was dissolved in DMF together with H-Asp(OBzl)-OtBu hydrochloride salt (41 mg, 130 µmol) and DIPEA (72 µL, 412 mmol). The mixture was cooled to 0°C and PyBOP (68 mg, 130 µmol) was added. The mixture was allowed to stir at room temperature for 4h (controlled by TLC). The reaction was quenched with water and the mixture was evaporated *in vacuo*. The crude product was purified by flash chromatography on silica $SiO_2$ gel with gradual eluting with DCM/methanol from 98:2 to 90:10 v:v%. Yield: 75 mg (68%) as a dark red solid. $^1H$ NMR (400 MHz,

$CDCl_3$) δ 8.37 (t, *J* = 13.4 Hz, 1H), 8.11 (d, *J* = 7.7 Hz, 1H), 7.80 (d, *J* = 8.3 Hz, 0H), 7.68 (d, *J* = 8.2 Hz, 0H), 7.41 – 7.16 (m, 17H), 7.00 (d, *J* = 13.4 Hz, 1H), 6.85 (d, *J* = 13.3 Hz, 1H), 6.59 (t, *J* = 12.5 Hz, 0H), 5.11 (dd, *J* = 11.2, 4.7 Hz, 2H), 4.77 (dt, *J* = 7.7, 6.1 Hz, 1H), 4.28 (dd, *J* = 10.5, 6.2 Hz, 2H), 4.20 – 4.09 (m, 3H), 3.69 – 3.58 (m, 2H), 3.28 (t, *J* = 7.5 Hz, 2H), 2.96 – 2.82 (m, 5H), 2.82 – 2.60 (m, 4H), 2.11 (q, *J* = 11.6 Hz, 8H), 1.68 (d, *J* = 4.4 Hz, 14H), 1.38 (s, 9H), 1.26 (dq, *J* = 14.3, 7.1 Hz, 2H), 0.87 (t, *J* = 7.3 Hz, 3H). $^{13}C$ NMR (101 MHz, $CDCl_3$) δ 174.01, 173.55, 172.97, 172.39, 170.49, 169.79, 151.05, 141.89, 141.86, 140.53, 140.35, 135.79, 129.06, 128.98, 128.47, 128.38, 128.30, 128.12, 125.38, 125.18, 121.92, 121.80, 111.77, 111.24, 104.63, 103.84, 81.90, 66.57, 49.79, 49.06, 48.96, 47.81, 45.66, 44.09, 43.67, 36.57, 32.73, 30.67, 30.07, 28.19, 28.06, 27.86, 27.80, 25.34, 24.27, 23.83, 20.19, 13.89. HRMS (ESI), m/z: [M+H]+ calcd for $C_{53}H_{70}N_4O_9S$, 938.4864; found, 938.4831.

**3-(4-(2-((E)-3-((Z)-1-(4-((1-(tert-butoxy)-3-carboxy-1-oxopropan-2-yl)amino)-4-oxobutyl)-3,3-dimethylindolin-2-ylidene)prop-1-en-1-yl)-3,3-dimethyl-3H-indol-1-ium-1-yl)-N-butylbutanamido)propane-1-sulfonate (13)**

Compound **12** (35 mg, 37 µmol) was dissolved in a mixture of DCM/EtOH (50:50) and Pd/C (2.7 mg, 2.6 µmol) wad added. The mixture stirred under $H_2$ at RT for 6h, filtered under Celite; and evaporated *in vacuo*. The crude product was used in the next step without further purification Yield: 31 mg (96%) as a dark red solid. $^{1}H$ NMR (400 MHz, MeOD) δ 8.45 (t, *J* = 13.4 Hz, 1H), 7.43 (d, *J* = 7.3 Hz, 2H), 7.34 (dd, *J* = 11.9, 3.8 Hz, 4H), 7.20 (dtd, *J* = 8.5, 5.0, 2.1 Hz, 2H), 6.55 – 6.39 (m, 2H), 4.56 (td, *J* = 6.0, 3.3 Hz, 1H), 4.12 (dq, *J* = 12.1, 5.6 Hz, 4H), 3.55 – 3.43 (m, 2H), 3.29 – 3.21 (m, 4H), 2.79 – 2.68 (m, 2H), 2.63 – 2.57 (m, 2H), 2.49 – 2.36 (m, 2H), 1.98 (dt, *J* = 34.4, 7.3 Hz, 6H), 1.67 (d, *J* = 1.9 Hz, 12H), 1.36 (s, 9H), 1.30 – 1.11 (m, 4H), 0.83 (td, *J* = 7.4, 5.4 Hz, 3H). $^{13}C$ NMR (101 MHz, MeOD) δ 174.69 (d, *J* = 11.8 Hz), 173.09, 172.50, 170.73 (d, *J* = 4.9 Hz), 150.95 (d, *J* = 11.8 Hz), 142.87 – 141.82 (m), 140.81 (d, *J* = 4.5 Hz), 128.65, 125.85 – 124.89 (m), 123.90 (d, *J* = 3.6 Hz), 122.08, 117.28, 111.25, 102.63, 81.53, 56.93, 49.27, 47.41, 47.20, 46.98, 43.18, 31.81 (d, *J* = 11.6 Hz), 30.53, 29.58, 28.83, 26.96, 26.87, 24.28, 22.86, 22.96 – 20.99 (m), 16.99, 12.85 (d, *J* = 3.5 Hz). HRMS (ESI), m/z: [M+H]+ calcd for $C_{46}H_{64}N_4O_9S$, 848.4394; found, 848.4392.

**3-(4-(2-((E)-3-((Z)-1-(6-(tert-butoxycarbonyl)-4,8,22-trioxo-26-(2-oxohexahydro-1H-thieno[3,4-d]imidazol-4-yl)-12,15,18-trioxa-5,9,21-triazahexacosyl)-3,3-dimethylindolin-2-ylidene)prop-1-en-1-yl)-3,3-dimethyl-3H-indol-1-ium-1-yl)-N-butylbutanamido)propane-1-sulfonate (14)**

Compound **13** (31 mg, 36.5 µmol) was dissolved in DMF together with Biotin-PEG-3-$NH_2$ (22 mg, 41.3 µmol) and DIPEA (26 µL, 146 µmol). The mixture was cooled to 0°C and HATU (15.5 mg, 41.3 µmol) and HOBt (2.5 mg, 20.6 µmol) were added. The mixture was allowed to stir at room temperature for 6h (controlled by TLC). The reaction was quenched with water and the mixture was evaporated *in vacuo*. The crude product was purified by flash chromatography on silica $SiO_2$ gel with gradual eluting with DCM/methanol from 98:2 to 90:12 v:v%. Yield: 27 mg (59%) as a dark red solid. $^{1}H$ NMR (400 MHz, MeOD) δ 8.45 (t, *J* = 13.4 Hz, 1H), 7.48 – 7.39 (m, 2H), 7.38 – 7.29 (m, 4H), 7.27 – 7.16 (m, 2H), 6.57 – 6.40 (m, 2H), 4.60 (ddd, *J* = 9.8, 7.4, 5.6 Hz, 1H), 4.38 (dd, *J* = 7.9, 4.8 Hz, 1H), 4.20 (dd, *J* = 7.9, 4.5 Hz, 1H), 4.17 – 4.05 (m, 4H), 3.58 – 3.46 (m, 10H), 3.43 (t, *J* = 5.6 Hz, 4H), 3.18 – 3.05 (m, 2H), 2.86 – 2.69 (m, 3H), 2.68 – 2.57 (m, 4H), 2.55 – 2.35 (m, 3H), 2.17 – 2.07 (m, 2H), 2.06 – 1.89 (m, 6H), 1.67 (d, *J* = 1.6 Hz, 11H), 1.62 – 1.39 (m, 3H), 1.36 (d, *J* = 1.5 Hz, 9H), 1.30 – 1.11 (m, 6H), 0.83 (td, *J* = 7.4, 2.3 Hz, 3H). $^{13}C$ NMR (101 MHz, MeOD) δ 174.78, 174.67, 174.60, 173.14, 173.05, 172.53, 172.18, 170.65, 170.61, 170.37, 164.64, 151.02, 150.87, 142.00, 141.92, 140.85,

140.82, 128.67, 128.62, 125.37, 125.32, 122.08, 111.29, 111.21, 102.65, 102.49, 81.70, 70.19, 70.06, 69.84, 69.20, 69.11, 61.96, 60.23, 55.61, 54.68, 54.46, 50.31, 49.29, 49.26, 49.24, 48.93, 46.62, 45.65, 44.95, 43.49, 43.18, 42.41, 39.68, 39.07, 38.94, 36.97, 35.35, 31.90, 31.72, 30.55, 29.67, 29.61, 28.78, 28.37, 28.16, 28.11, 26.98, 26.91, 25.45, 24.37, 23.27, 22.99, 22.91, 22.71, 22.28, 19.85, 19.73, 12.89, 12.86, 11.77. HRMS (ESI), m/z: [M+H]+ calcd for $C_{64}H_{96}N_8O_{13}S_2$, 1248.6538; found, 1248.6518.

**3-(N-butyl-4-(2-((E)-3-((Z)-1-(6-carboxy-4,8,22-trioxo-26-(2-oxohexahydro-1H-thieno[3,4-d]imidazol-4-yl)-12,15,18-trioxa-5,9,21-triazahexacosyl)-3,3-dimethylindolin-2-ylidene)prop-1-en-1-yl)-3,3-dimethyl-3H-indol-1-ium-1-yl)butanamido)propane-1-sulfonate (15)**

Compound **14** was dissolved in a mixture of DCM/TFA (70:30) and the reaction was allowed to stir 4h at RT. The reaction mixture was evaporated *in vacuo* and dried under high vacuum overnight. The crude product was used in the next step without further purification. The quantity of product obtained was too small to record an NMR spectrum. The product was characterized by HRMS (ESI), m/z: [M-$H_2O$]+ calcd for $C_{60}H_{88}N_8O_{13}S_2$, 1174.575; found, 1174.577.

**NHS ester preparation**

A corresponding cyanine carboxylic acid derivative (**7**, **8**, **9** and **10**) was dissolved in dry DCM (1 mL) followed by DIPEA (1.5 eq) and TSTU (1.5 eq). The reaction was maintained by stirring at room temperature for 6h. The reaction was quenched with water, DCM was added and the organic layer was washed with aqueous 5% citric acid and brine. Then, it was dried over $Na_2SO_4$, filtrated and concentrated under vacuum. NHS esters of cyanine derivatives (MemGraft-Cy3, MemGraft-Cy5 and **1**, respectively) were obtained as a fine powder and used in microscopy without any intermediate purification.

**Cell Lines, Culture Conditions, and Treatment**

U87 (ATCC HTB-14) cells and Hela (ATCC CCL-2) cells were grown in Eagle's Minimum essential medium (EMEM, Gibco Invitrogen), supplemented with 10% fetal bovine serum (FBS, Lonza), 2 mM L-glutamine (Gibco-Invitrogen), 1% non-essential amino acid solution (Gibco-Invitrogen) and sodium pyruvate 1 mmol/L at 37 °C in a humidified 5% $CO_2$ atmosphere. Cells were seeded onto a chambered coverglass (IBIDI) at a density of 5 × $10^4$ cells/well 24 h before the microscopy measurement.

For microscopy imaging, the attached live cells in IBIDI dishes were washed once with warm Hanks' balanced salt solution (HBSS, Gibco- Invitrogen); after that, 1 mL of a corresponding dye solution in HBSS was added and the cells were incubated for 5 min at room temperature. Then, the attached cells in IBIDI dishes were washed twice with HBSS before imaging. Serum resistance experiments were performed with the same protocol followed by the incubation of the dye with 20% FBS in PBS. For fixation and permeabilization experiments, the dye solution was removed from the cells and the attached cells were washed twice with HBSS and incubated for 15 min with 4% (w/v) formaldehyde solution in phosphate-buffered saline (PBS) at room temperature. The fixative solution was removed and the cells were washed by pipetting PBS against the side of the dish 3 times. Then, the cells were incubated with 1 mL of 0.1% Tween-20 in PBS for 12 min at room temperature. The permeabilization solution was removed and washed 3 times with PBS.

For co-culture experiments, the cells were seeded in a 6-well plate at a density of 5 × $10^4$ cells/well 24 h before the experiment. The attached cells were stained by MemGraft-Cy3 or MemGraft-Cy5 as described above, washed twice with HBSS, detached with StemPro Accutase by incubating for 10 min at 37°C, centrifuged, and distributed in IBIDI dishes filled with the culture medium described previously. After 5 h or 29 h of incubation, the co-cultured cells were imaged in the culture medium without more treatment.

In the experiments on blocking amino-group sites membrane proteins, the U87 cells were treated with Pierce™ Sulfo-NHS-Acetate at 1 mM in HBSS for 60 min at 37°C. The blocking solution was removed and the cells were washed with PBS and stained with the corresponding dye solution following the previously described protocol.

**Fluorescence Microscopy**

Confocal imaging of cells was performed on a Leica TCS SP8 confocal microscope with a HCX PL APO 63×/ 1.40 OIL CS2 objective and two 12-bit photomultipliers. The excitation light was provided by lasers of 488 nm, 552 nm, and 638 nm for MemBright-488, MemGraft-Cy3, and MemGraft-Cy5, respectively. The fluorescence from these three probes was detected at the corresponding spectral ranges: 500−540 nm, 570−620 nm, and 650-700 nm. All the parameters at each channel were left constant; the illumination power was adjusted to achieve a good signal for each probe. When the performances of the probes were compared, all instrumental conditions were fixed.  All the images were processed using ImageJ software.

Video microscopy experiments on live cells were performed on a Leica DMIRE 2 microscope with a HXC PL APO 40x/ 1.25 oil objective equipped with a Photometric Prime camera. The sample chamber is controlled by a thermostatic chamber and $CO_2$-controlled incubation chamber for long-term live imaging conditions. The system is piloted by the software Metamorph version 7.8.13.0 from Molecular Devices. The excitation light is provided by the LED light source Lumencor SOLA III equipped for exciting light in 5 different channels (DAPI, GFP, Cy3, Cy5 and transmitted light. The images were processed and converted into videos using ImageJ software.

**Flow Cytometry Analysis**

U87 cells were seeded at a density in 1×$10^6$/well in a 6-well plate 24h before flow cytometry measurement. For cytometry analysis, the attached live cells were washed once with warm Hanks' balanced salt solution (HBSS, Gibco-Invitrogen). After that, 5 mL of a corresponding dye (MemGraft-Cy3 or disulfo-Cy3-NHS) solution at 0.1 and 1 µM in HBSS was added and the cells were incubated for 10 min at room temperature. Then, the attached cells in IBIDI dishes were washed twice with HBSS. The cells were then detached with StemPro Accutase incubation for 10 min at 37°C, centrifuged, and resuspended in a solution of BSA 1% in PBS. This suspension was passed through a 50 µm nylon mesh filter. For the experiment, the cells were analyzed using a MACSQuant® VYB flow cytometer. MemGraft-Cy3-NHS and commercial disulfoCy3-NHS were excited with a 561 nm laser and the fluorescence was collected at 579-593nm (PE and SYTOX orange filter, named Y1). The detectors were calibrated in by the light scattering and fluorescence using MACSQuant® fluorescent calibration beads.

**Cell viability assay**

The MTT assay was performed to calculate the toxicity of the probe in mammalian cells. Five thousand HeLa cells were seeded per well and grown in 96-well plates. After 24 h of seeding, the cells were treated with different concentrations (0.02, 0.1, 0.5, 1, 2, and 5 µM) of the probes, respectively, and incubated for 15 min. After that, the staining solution was replaced with fresh medium and the cells were left for 24h at 37°C. At the end of this incubation, the medium was aspirated from all the wells, and the cells were incubated further in fresh growth medium with 0.5 mg $mL^{-1}$ of MTT for 4 h. The medium containing MTT was then removed, after which DMSO was added to solubilize the formazan. The absorbance values were recorded using a TECAN Spark plate reader at 570 nm. The cell viability was calculated as the absorbance with respect to the positive control as a reference, in which the cells were treated with the solvent (DMSO) alone. In the negative control used as a cytotoxicity reference, cells were treated with TX-100 1% for 1h. The results presented here are a mean of 8 experiments ± standard deviation.

**Model membrane conjugation assay**

Large Unilamellar Vesicles (LUVs) were prepared by the following procedure. A stock solution of the corresponding lipids (DOPC and DOPE, 1/1, mol/mol) in chloroform was placed into a round-neck flask, after which the solvent was evaporated in vacuo. PBS was added so that the final lipid concentration was 1 mM and the mixture was sonicated. The obtained suspension of multilamellar vesicles was extruded using a Lipex Biomembranes extruder (Vancouver, Canada). The size of the filter was first 0.2 µm (7 passages) and thereafter 0.1 µm (10 passages). This generates monodisperse LUVs with a mean diameter of 0.11 µm as measured by dynamic light scattering method with a Malvern Zetasizer ZSP (Malvern Instruments S.A.). LUVs were labeled by the addition of the DMSO stock solution of the MemGraft-Cy3 probe to a final concentration of 250 µM in PBS, and incubation with LUVs for 20 min using an Eppendorf Thermomixer© C, with a rotation of 800 rpm at room temperature. The labeled liposomes were separated from the unreacted probe using a disposable size exclusion chromatography column, Illustra™, NAP™-5 from Cytiva and the fraction containing LUVs was analyzed using a MALDI-TOF MS system (Microflex ® LRF, Bruker). The optimal conditions for observation of phospholipid derivatives were found as follows: matrix DCTB in chloroform in Reflector Negative ion mode.

**Protein extraction**

U87 cells were seeded at a density in $5\times10^6$/well in a petri dish 24h before the experiment. The attached live cells were washed once with PBS. After that, 5 mL of a corresponding dye (MemGraft-Cy3-NHS or MemGraft-Cy3-Mal) solution at 1 µM in HBSS was added and the cells were incubated for 10 min at room temperature. Then, the attached cells in dishes were washed with PBS and put in growth media. The cells were resuspended in the growth media by scraping the cells off the surface of the plate with a cell scraper. The cell suspension was centrifuged at 300 × g for 5 minutes.

The cell pellet was washed with 3 mL of Cell Wash solution from the Mem-PER$^{TM}$ Plus Membrane Protein Extraction Kit from ThermoFisher, and centrifuged at 300 x g for 5 minutes. The supernatant was removed and the cells were resuspended in 1.5 mL of Cell Wash solution and transferred to a 2 mL centrifuge tube. The cells were centrifuged at 300 x g for 5 minutes and the supernatant was removed. 0.75 mL of Permeabilization Buffer from the same kit were added to the cell pellet and the mixture was vortexed. The suspension was incubated 10 minutes at 4°C under constant mixing while permeabilization occurs. The permeabilized cells were centrifuged for 15 minutes at 16.000 x g to separate cytosolic proteins. The supernatant with cytosolic proteins was entirely transferred to a new tube and stored in the permeabilization buffer. 0.5 mL of Solubilization Buffer from the same kit was added to the pellet and the cells were resuspended by pipetting up and down, and incubated at 4°C for 30 minutes with constant mixing. The suspension was then centrifuged at 16.000 x g for 15 minutes at 4°C. The supernatant containing membrane proteins and solubilized membranes was transferred to a new tube.

**Protein concentration assay**

Protein contents of extracted protein samples from MemGraft treated U87 cells were checked using colorimetric BioRad DC protein Assay based on Lowry method following supplier's microplate assay protocol. The tests have been performed in transparent polystyrene 96-wells microplates without lid and absorbance measurements at 750 nm done using a Tecan Spark microplate reader. A standard curve has been established using BioRad quick start BSA 7 standards set from 0.125 to 2.0 mg/mL. Typical protein concentrations obtained in extracted samples were close to 0.5 µg/mL.

**SDS-Page**

Purified membrane proteins in solubilization buffer and cytosolic proteins in permeabilization buffer were directly diluted with 4x Laemmli sample buffer (BioRad) and gels (BioRad Criterion 4-20%, 1mm TGX 18-wells precast Gels) were loaded with 30 µL/wells. BioRad Precision Plus Protein™ All Blue Prestained Protein Standards (10-250 kD) was used as ladder. Resulting gels were run in 1x Tris-Glycine Buffer pH = 9 (Euromedex) with 0.2 % SDS added with BioRad Powerpac generator (150 V, 300 W) for about 35 min. After SDS-Page migration, a fluorescence image of the gel were took using Syngen G:Box Chemi XRQ (Ex: Green LED 520-550 nm Em: 605 nm filter). After that, gels were stained with Coomassie solution overnight and then destained using successive wash in 1 % Acetic acid baths until non-specific staining was completely removed.

**Magnetic cell manipulation**

U87 cells were seeded at a density in $1\times10^6$ / well 24h before the experiment. The attached live cells were washed once with PBS. After that, 2 mL of MemGraft-Cy3-Biotin solution at 1 µM in HBSS was added in 3 wells and MemGraft-Cy5-NHS solution at 1 µM in HBSS was added in 3 other wells and were incubated for 10 min at room temperature. Then, all the attached stained cells in dishes were washed with PBS and detached by 5 min incubation at 37°C with an Accutase solution and the cells with same staining were mixed together. The suspensions of cells were centrifuged for 5 minutes at 1.500 rpm. The supernatant was discarded and the cells were resuspended in OptiMEM. The two batches of cells stained with MemGraft-Cy5-NHS and MemGraft-Cy3-Biotin were mixed together by pipetting up-and-down. One IBIDI dish was reseeded in OptiMEM with this suspension mixture as a negative control. The rest of the suspension was mixed with 3 times pre-washed (because of toxic azide in the formulation) Pierce™ Streptavidin Magnetic Beads from Thermo Fisher, distributed in 0.5 mL centrifuge tubes and placed on a magnetic separation stand from Promega. After 2 minutes, the supernatant containing a part of the cells was carefully collected and the cells were seeded in a new IBIDI dish in OptiMEM. The rest of the cells were taken out from the magnetic separation stand, suspended in OptiMEM, and reseeded in a new IBIDI dish. The 3 dishes were incubated for adhesion for 3 h at 37°C before the imaging.

### Additional microscopy images

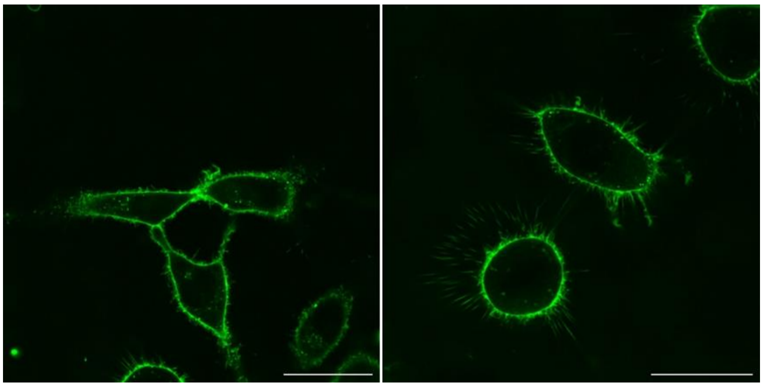

**Figure S2.** Fluorescence imaging of HeLa cells labelled with MemGraft-Cy3. Dye concentration was 500 nM. Scale bar: 30 µm.

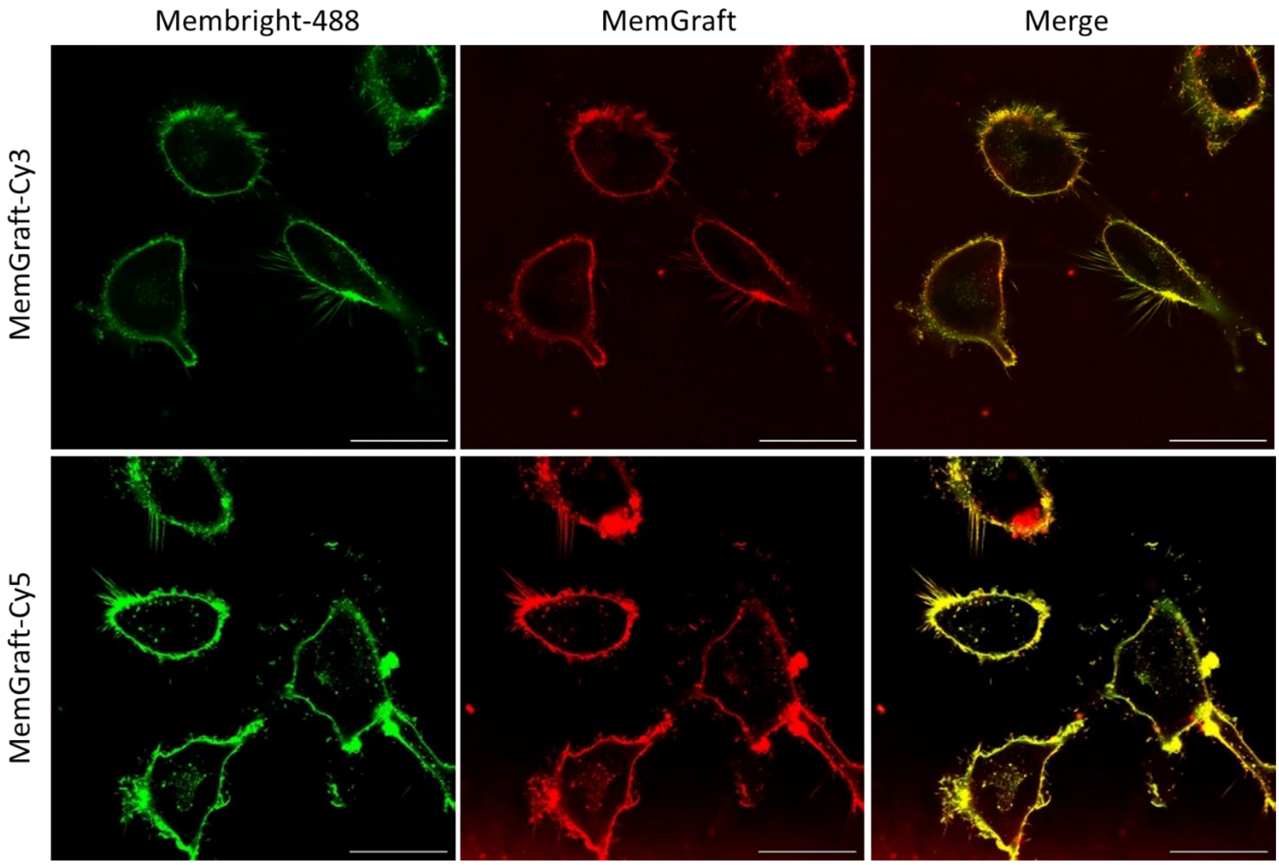


**Figure S3.** Fluorescence imaging of U87 cells labelled with MemGraft-Cy3 and MemGraft-Cy5 in comparison to MemBright-488. Left panels: MemBright-488, middle panels: MemGraft-Cy3 and MemGraft-Cy5; right panels: merged images. Colocalization Pearson's coefficients are 0.84 and 0.85 for MemGraft-Cy3 and MemGraft-Cy5, respectively. Dye concentrations: 500 and 200 nM for MemGraft and MemBright-488, respectively. Scale bar: 30 µm.

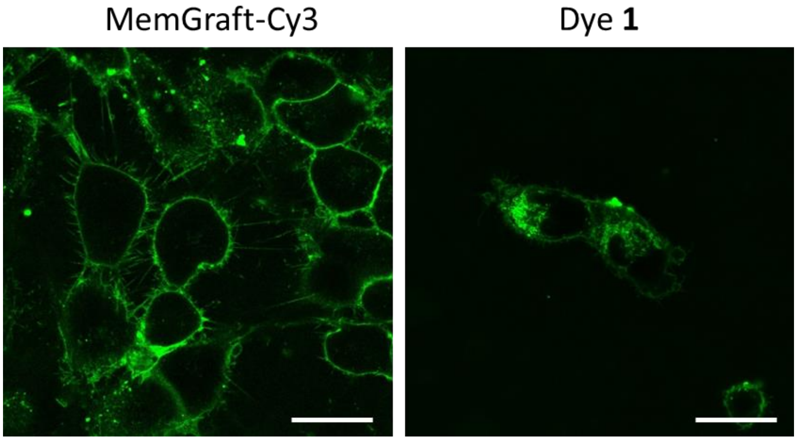


**Figure S4.** Comparison of MemGraft-Cy3 dye with a control dye **1** without butyl anchor at 100 nM concentration. Scale bar: 30 µm.

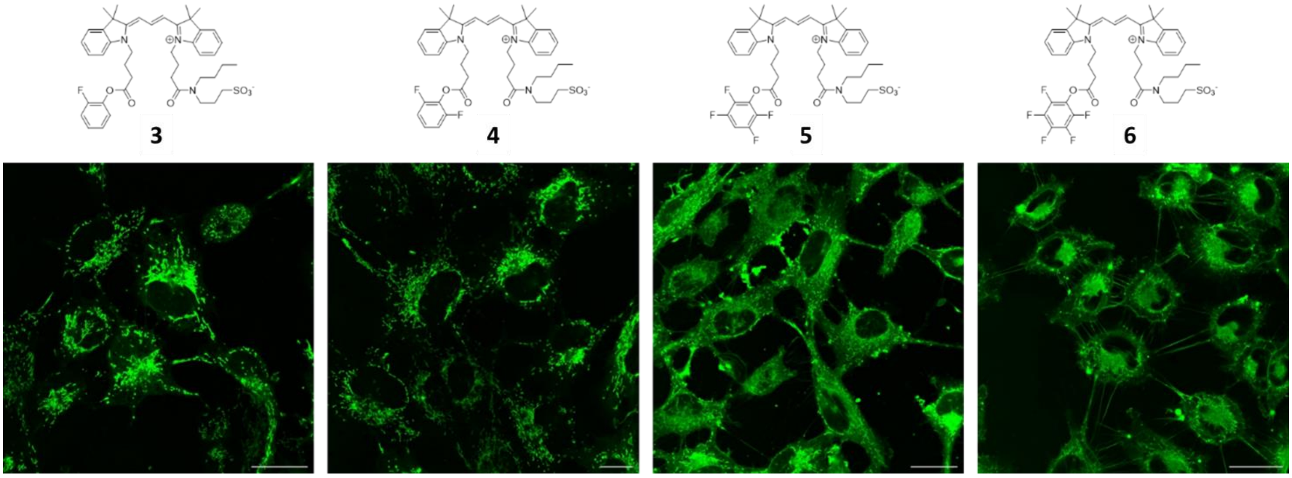


**Figure S5.** Fluorescence labelling of U87 cells with dyes **3**-**6** after 15 min incubation and washing. Dye concentrations: 500 nM. Scale bar: 30 µm.

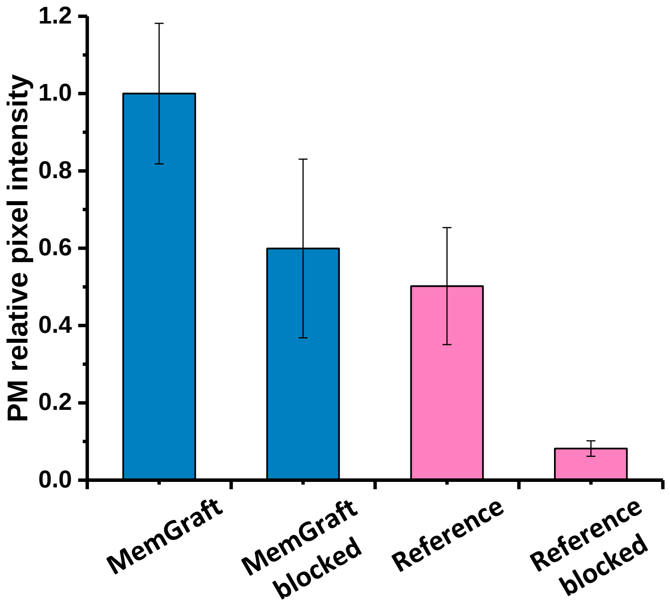


**Figure S6.** Blocking of PM labelling by pretreatment with Sulfo-NHS-acetate. Fluorescence intensity of cells labelled with MemGraft-Cy3 and a reference dye (disulfo-Cy3-NHS) without and with prior incubation with 1 mM of Sulfo-NHS-Acetate for 60 min at 37°C. The concentration of the dyes was 500 nM. The incubation time was 5 min. The error is s.d.m. based on ~20 cells.

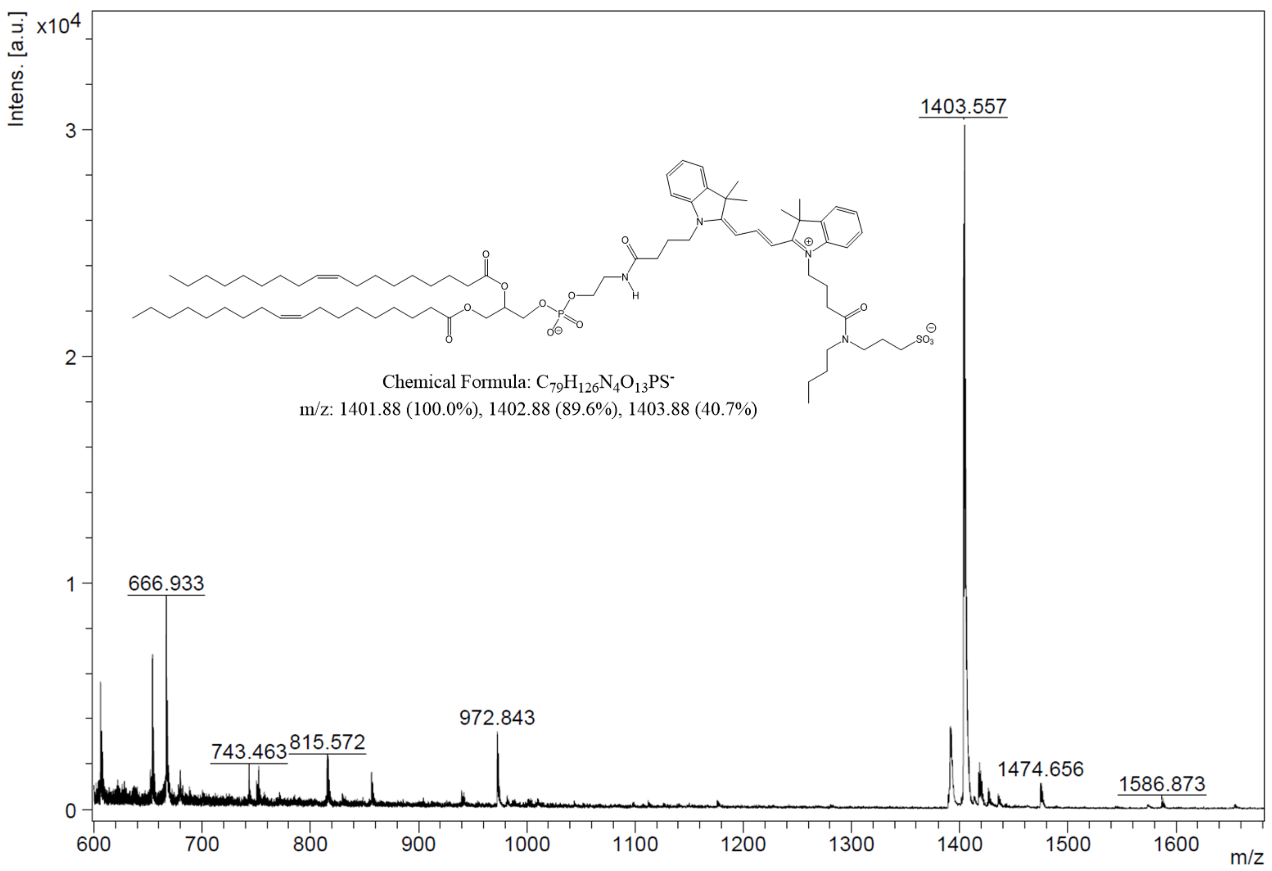


**Figure S7.** Mass analysis of expected conjugate of MemGraft-Cy3 with DOPE lipid after reaction of the probe with liposomes composed of DOPC/DOPE mixture, 1/1, mol/mol.

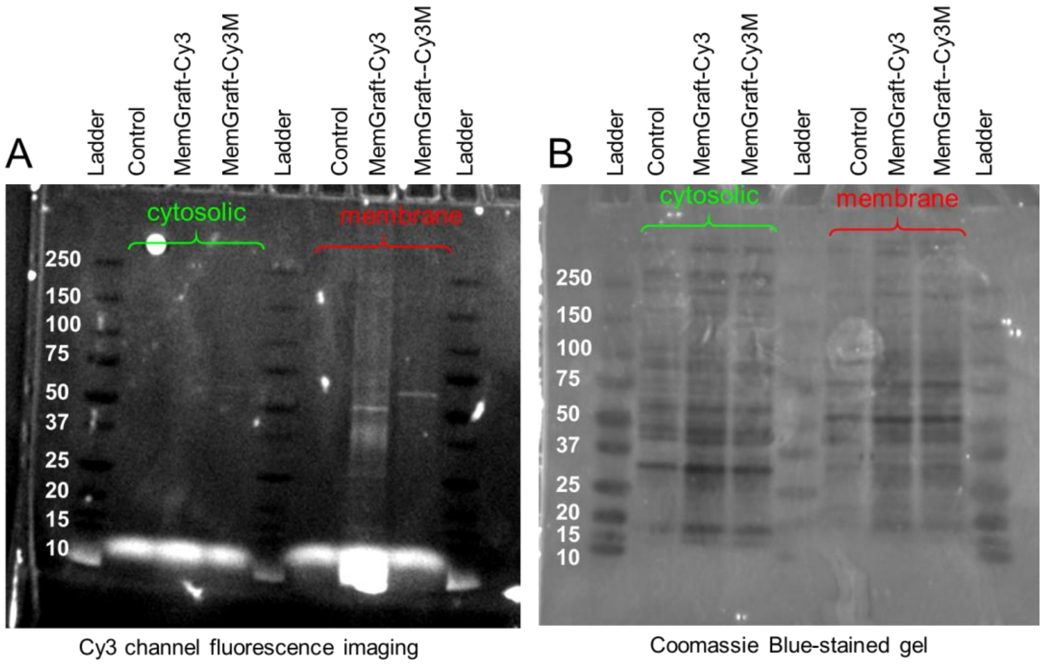


**Figure S8.** SDS-polyacrylamide gel electrophoresis (SDS-PAGE) results of cytosolic and membrane proteins extracted from U87 cells stained with MemGraft-Cy3, MemGraft-Cy3M or without staining (control). (A) fluorescence imaging of the gel with Cy3 channel. (B) Brightfield image of the Coomassie Blue-stained gel.

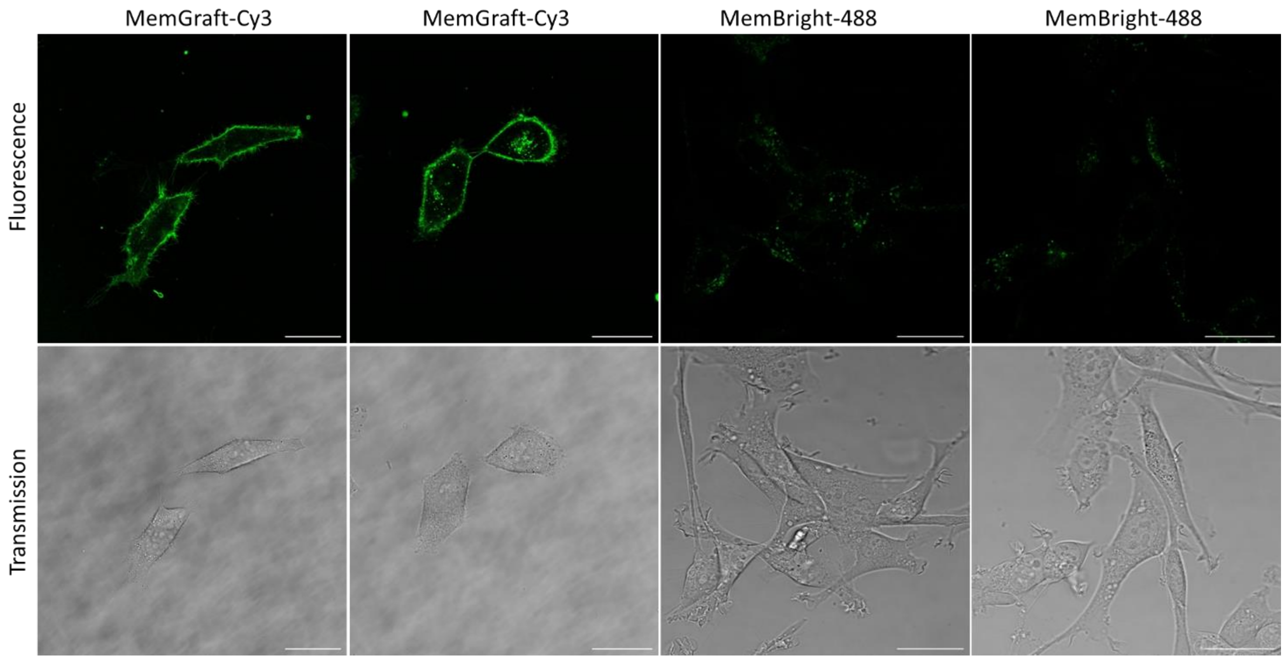


**Figure S9.** Fluorescence imaging of U87 cells labelled with MemGraft-Cy3 and MemBright-488 after trypsinization of seeding to a new microscopy plate and incubation for 24h in the full growth medium with 10% FBS (serum). Concentration of MemGraft-Cy3 and MemBright-488 were 1 and 0.2 µM, respectively. The results are shown in duplicates for each condition. Scale bar: 30 µm.

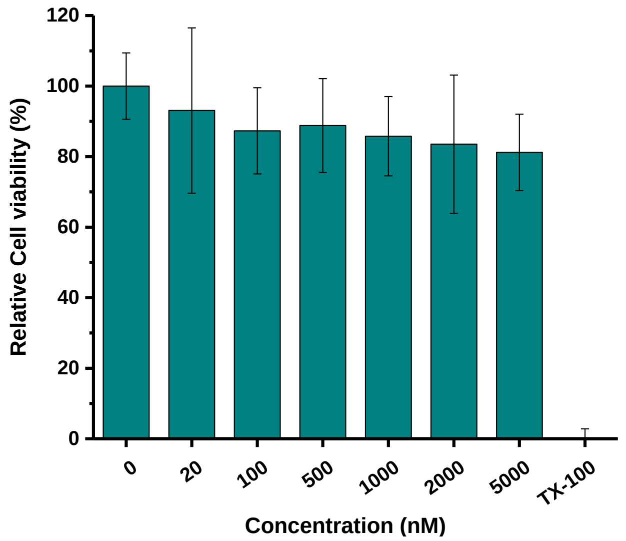


**Figure S10.** Viability of Hela cells after incubation with MemGraft-Cy5 at different concentrations for 24h. Data for MemGraft-Cy3 are not available because of potential cross-talk of the dye with MTT assay.

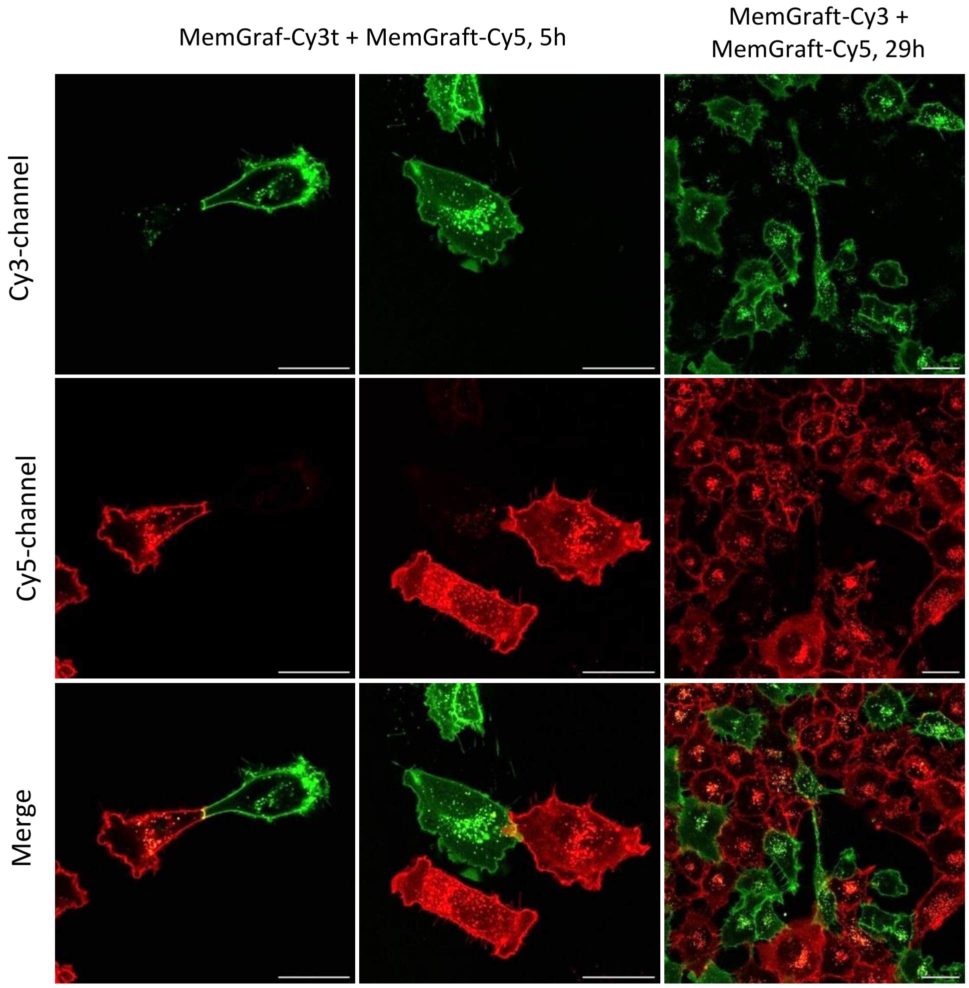


**Figure S11.** Additional examples of co-seeding of cells stained with MemGraft-Cy3 and MemGraft-Cy5 (C): after 5h ; (D): after 29h. Cells stained only with MemGraft-Cy3 or MemGraft-Cy5 are shown respectively in A and B Scale bar: 30 μm.

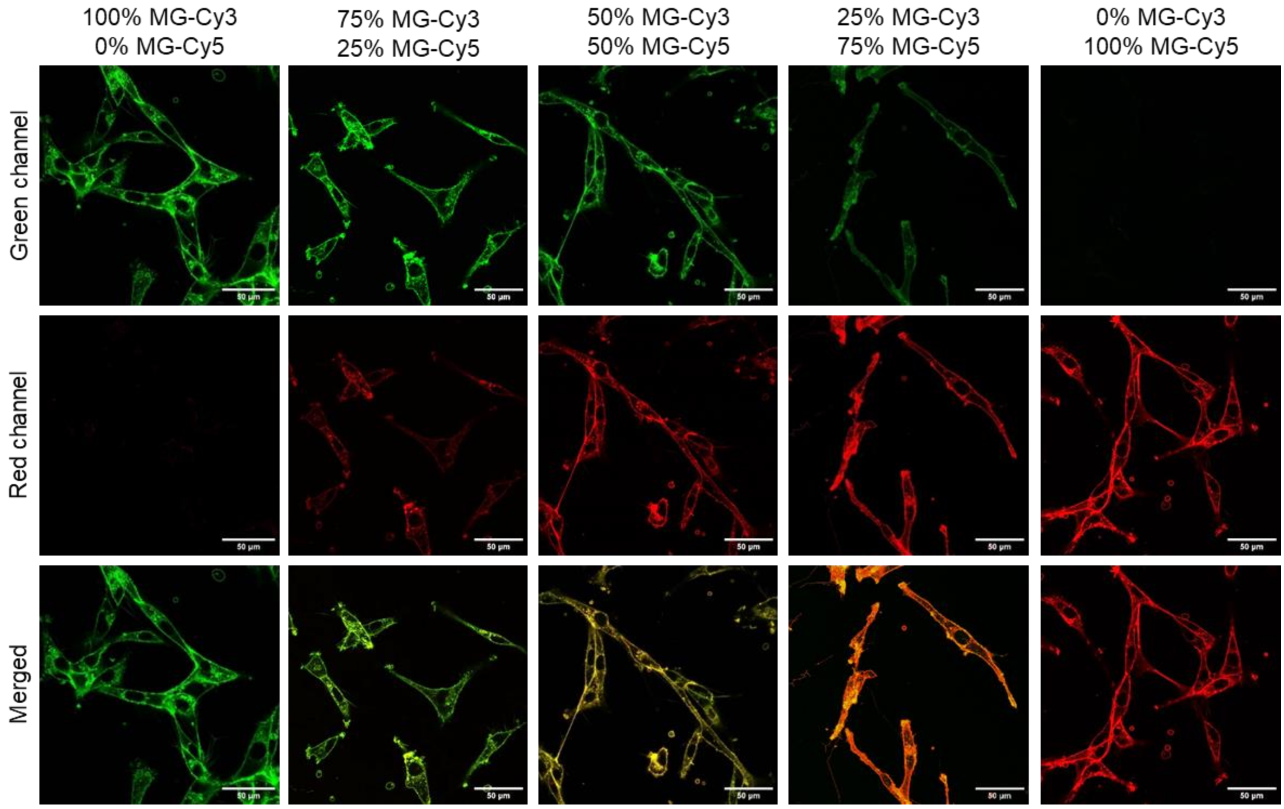


**Figure S12.** Confocal fluorescence microscopy of U87 cells stained with varied molar % of MemGraft-Cy3 (MG-Cy3) / MemGraft-Cy5 (MG-Cy5): 100%/0%, 75%/25%, 50%/50%, 25%/75%, 0%/100% (left to right). Upper panels: green channel (Cy3). Middle panels: red channel (Cy5). Lower panels: Merged channels. Total dye concentration was 1 µm in each case. Scale bars: 100 µm.

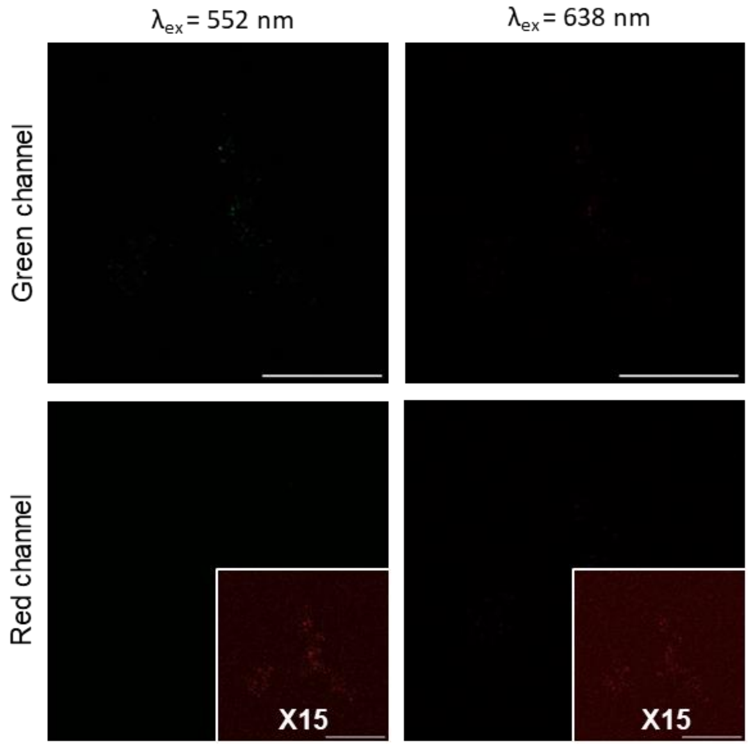


**Figure S13.** Confocal fluorescence microscopy of U87 cells incubated for 30 min with streptavidin-Cy5 adduct only as a control. On the left panel, the images are recorded with excitation of Cy3 (552 nm). On the right panel, the images are recorded with excitation of Cy5 (638 nm). Upper panels: green channel (Cy3). Lower panels: red channel (Cy5). Final streptavidin-Cy5 concentration was 0.1 mg/mL Scale bars: 30 μm.

**Video S1.** Fluorescence video imaging of U87 cells labelled with MemGraft-Cy3. Recording time was 3 hours with 1 frame / min. Dye concentration was 1 μM. Scale bar: 50 μm

**Video S2.** Fluorescence video imaging of co-seeded U87 cells labelled with MemGraft-Cy3 and MemGraft-Cy5. Recording time was 1 hour with 1 frame / min. Dye concentration was 1 μM. Scale bar: 50 μm
